# Dynamics of sister chromatid resolution during cell cycle progression

**DOI:** 10.1101/283770

**Authors:** Rugile Stanyte, Johannes Nuebler, Claudia Blaukopf, Rudolf Hoefler, Roman Stocsits, Jan-Michael Peters, Daniel W. Gerlich

## Abstract

Faithful genome transmission in dividing cells requires that the two copies of each chromosome’s DNA package into separate, but physically linked, sister chromatids. The linkage between sister chromatids is mediated by cohesin, yet where sister chromatids are linked and how they resolve during cell cycle progression has remained unclear. Here, we investigated sister chromatid organization in live human cells using dCas9-mEGFP labelling of endogenous genomic loci. We detected substantial sister locus separation during G2 phase, irrespective of the proximity to cohesin enrichment sites. Almost all sister loci separated within a few hours after their respective replication, and then rapidly equilibrated their average distances within dynamic chromatin polymers. Our findings explain why the topology of sister chromatid resolution in G2 largely reflects the DNA replication program. Further, these data suggest that cohesin enrichment sites are not persistent cohesive sites in human cells. Rather, cohesion might occur at variable genomic positions within the cell population.

## Introduction

To transmit the genetic information through generations, cells must duplicate each chromosome’s DNA and package both copies into separate cytological bodies, termed mitotic sister chromatids. In vertebrate cells, the replicated DNA of each chromosome initially co-localizes within the same nuclear territory (Bickmore and van Steensel, 2013; Nagasaka et al., 2016). Sister chromatids become visible as separate rod-shaped structures only when cells enter mitosis, a few minutes before the nuclear envelope disassembles (Kireeva et al., 2004; Liang et al., 2015; Nagasaka et al., 2016). However, individual genomic sites labelled by fluorescence in situ hybridization (FISH) often appear as pairs of fluorescent foci after their replication, many hours before cells enter mitosis (Selig et al., 1992; Azuara et al., 2003; Mlynarczyk-Evans et al., 2006; Schmitz et al., 2007; Nishiyama et al., 2010). Hence, at least parts of replicated chromosomes move apart long before sister chromatids become visible as separate cytological bodies. How this is regulated in time and to what extent it is influenced by the genomic neighbourhood is unclear.

Although sister chromatids resolve during mitosis, they remain physically linked to enable correct attachment to the mitotic spindle (Nasmyth and Haering, 2009). This is mediated by the cohesin protein complex (Michaelis et al., 1997; Guacci et al., 1997), which forms a tripartite ring to topologically link DNA of sister chromatids (Gruber et al., 2003; Haering et al., 2008). Cohesin’s interaction with chromosomes is regulated throughout the cell cycle by various co-factors. Before DNA replication, cohesin binds to chromosomes with a short residence time (Gerlich et al., 2006; Ladurner et al., 2016; Hansen et al., 2017; Rhodes et al., 2017), whereby the protein WAPL promotes dynamic turnover (Kueng et al., 2006). During S-phase, a fraction of cohesin converts to a stably-chromatin-bound state (Gerlich et al., 2006) by acetylation of the SMC3 subunit and binding of Sororin (Schmitz et al., 2007; Ladurner et al., 2016). Sororin stabilizes cohesin on chromatin by counteracting WAPL; this function is required to maintain sister chromatid cohesion from S-phase until mitosis (Schmitz et al., 2007; Nishiyama et al., 2010; Ladurner et al., 2016).

Besides holding sister chromatids together, cohesin also organizes chromatin within sister chromatids. Chromatids contain domains with high contact probability, termed topologically associated domains (TADs) (Dixon et al., 2012; Nora et al., 2012). Cohesin enriches at the boundaries of TADs and is required for their formation (Rao et al., 2014; Schwarzer et al., 2016; Wutz et al., 2017; Gassler et al., 2017; Zuin et al., 2014a; Rao et al., 2017). It has been hypothesized that cohesin forms TADs by extruding chromatin loops, whereby the boundaries are specified by the protein CTCF (Nasmyth, 2001; Fudenberg et al., 2016; Sanborn et al., 2015; Rao et al., 2017; Busslinger et al., 2017).

Genomic sites enriched for cohesin might not only represent TAD boundaries but might also represent sites of preferential sister chromatid cohesion. In fission yeast, cohesin ChIP-sequencing peaks that co-localize with the cohesin loading factor Mis4 (NIPBL in humans) represent sites of persistent sister chromatid linkage (Bhardwaj et al., 2016). In human cells, however, there is very little overlap between cohesin ChIP-sequencing peaks and NIPBL (Zuin et al., 2014b). Moreover, cohesin can laterally diffuse along DNA (Davidson et al., 2016; Stigler et al., 2016) and relocate to distant genomic regions (Lengronne et al., 2004; Busslinger et al., 2017). Therefore, a substantial fraction of cohesin might occupy variable genomic positions in different cells within a heterogeneous population, which would not be detectable by conventional ChIP. Currently available chromosome conformation capture techniques also cannot detect linkage sites between sister chromatids because of the identical DNA sequences of sister chromatids. Whether the human genome encodes distinct sites of enhanced sister chromatid cohesion, and how these would relate to cohesin enrichment sites, have remained unknown.

In addition to cohesion, other factors might influence the spatial pattern of sister chromatid resolution. In vertebrate cells, the degree of sister locus separation during G2 appears to be elevated at sites with high transcriptional activity (Azuara et al., 2003; Mlynarczyk-Evans et al., 2006) and at genomic regions that replicate early (Selig et al., 1992). However, only few genomic loci have been investigated, and whether cohesin enrichment sites in the genomic vicinity affect sister locus resolution has remained unclear. Moreover, prior studies of sister locus separation relied on FISH, which might induce artefacts owing to harsh sample preparation procedures.

Progress in live-cell genome labelling methodology provides new opportunities to investigate the spatial distribution of sister chromatid linkage sites and to follow their resolution as cells progress from S-phase to mitosis. Here, we used CRISPR/Cas9 technology (Chen et al., 2013) to generate a collection of human cell lines with fluorescently labelled endogenous genomic loci and investigated the resolution of sister chromatids by live-cell microscopy. Our study reveals that sister chromatids resolve by a dynamic process that initiates long before mitotic entry.

## Results

### A human cell line collection with fluorescently labelled endogenous genomic loci

To systematically probe the topology of sister chromatids in living human cells, we used single guide RNA (sgRNA)/Cas9 technology to target a catalytically inactive Cas9-monomeric enhanced green fluorescent fusion protein (dCas9-mEGFP) to unique genomic regions. This technology efficiently labels unique repeat regions in the genome of human cells without compromising cell viability and with minimal effects on DNA damage (Chen et al., 2013). We identified candidate genomic sites for fluorescent labelling by analyzing the human Tandem Repeat Database (Gelfand et al., 2007) and mapping all genomic regions that contain at least 20 copies of a single sgRNA target site within 20 kb. We selected 113 regions, and individually introduced their corresponding sgRNAs into HeLa cells expressing dCas9-mEGFP. Imaging of live cells by confocal microscopy revealed discrete, nuclear foci for 47/113 regions. We pursued 16 regions that reside on 11 different chromosomes, are at least 0.5 Mbp from centromeres/telomeres, and are located within various genomic contexts, including protein-coding genes, long non-coding RNA genes, and intergenic regions (Table S1). To this end, we generated 16 HeLa cell lines that stably express a single sgRNA and doxycycline-inducible dCas9-mEGFP. After induction of dCas9-mEGFP, we imaged live cells by confocal microscopy and analyzed all cells expressing low to middle levels of dCas9-mEGFP (to avoid saturation of the signal by nucleoplasmic background fluorescence). For each cell line, we detected a small number of fluorescent foci in the nucleus of >90 % of the analyzed cells, whereby the number of foci per nucleus matched the corresponding number of sgRNA target alleles in HeLa cells (Fig. S1) (Landry et al., 2013). The mean fluorescence of different alleles within individual nuclei varied to some extent, which might be due to their different z-position relative to the focal sectioning planes, or which might reflect allele-specific copy number variations of the sgRNA target sequence. Overall, the high locus-labelling efficiency in our collection of 16 cell lines enabled us to assess the topology of sister chromatids in distinct genomic contexts.

Labelling of genomic sites with dCas9-mEGFP/sgRNA might perturb cell essential processes such as DNA replication. To test this, we investigated cell proliferation, mitotic duration, and nuclear morphology. We performed long-term time-lapse microscopy of eight randomly selected cell lines with dCas9-mEGFP/sgRNA labelled loci and found that their nuclear morphology and proliferation rates were indistinguishable from control cells, and that they progressed through mitosis without delay (Fig. S2, A-F). Hence, dCas9-mEGFP/sgRNA labelling does not impair essential cell functions.

### Monitoring sister chromatid resolution in live cells

To study the organization of sister chromatids in each cell line, we collected mitotic cells by mechanical shake-off, seeded them into chambered coverslips, and imaged them by 3D confocal time-lapse microscopy throughout an entire cell cycle (Fig. 1, A-D). For each cell, we monitored entry into the following mitosis, based on cell rounding and the release of the nucleoplasmic pool of dCas9-mEGFP into the cytoplasm (Fig. 1 B). We selected image frames from 2.4 – 0.6 h preceding mitosis, which corresponds to G2 phase in HeLa cells (Held et al., 2010), for further analysis.

**Figure 1.**
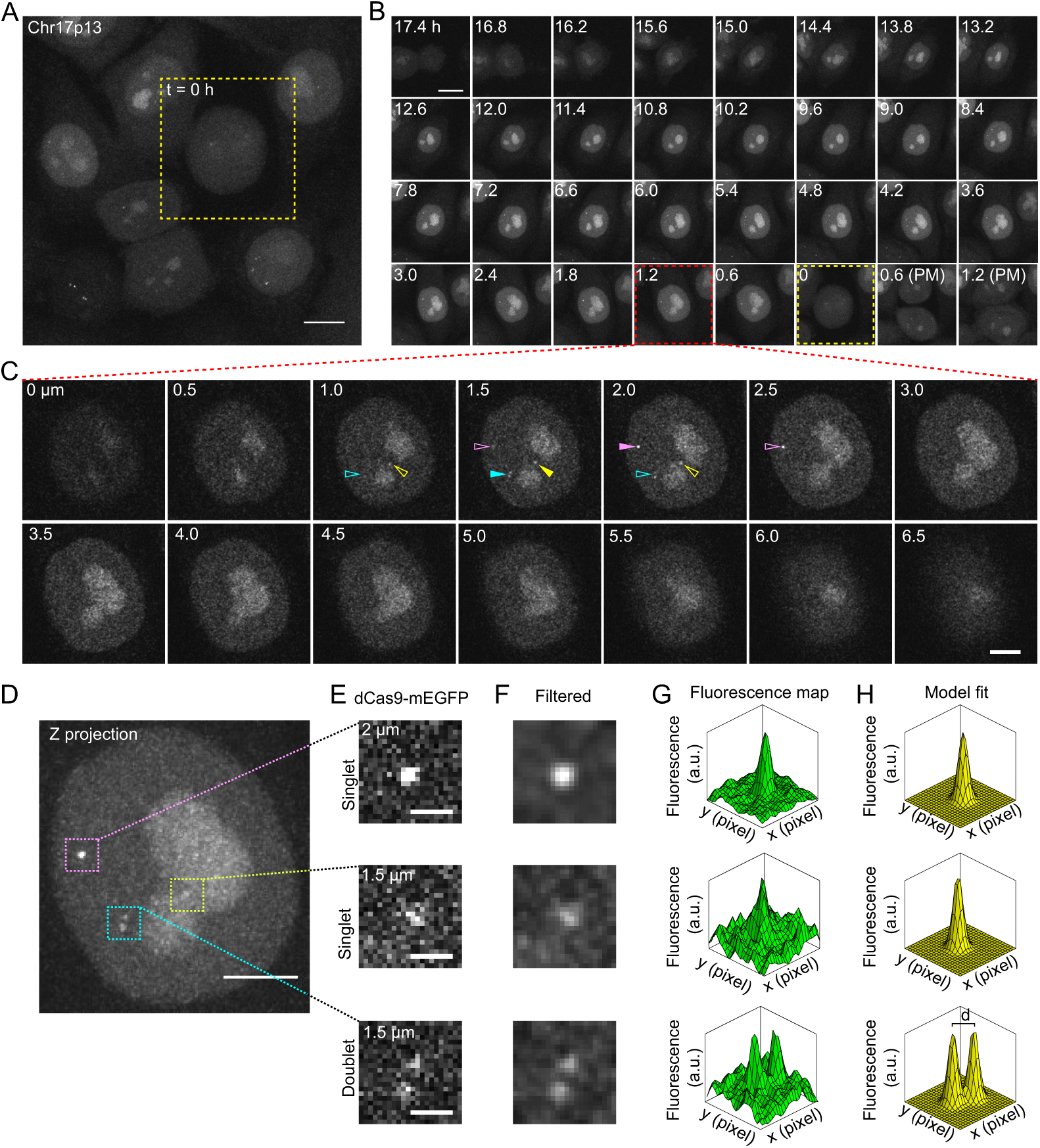
3D live-cell imaging of endogenous genomic loci throughout the cell cycle. (A-D) Confocal live-cell microscopy of HeLa cells expressing dCas9-mEGFP and sgRNA targeting Chr17p13 over 24 h with a time-lapse of 36 min and 21 z-sections at 0.5 μm spacing. (A) Maximum intensity projection at 17.4 h after imaging onset (yellow square indicates mitotic cell). (B) Time-lapse of maximum intensity projections for cell shown in (A); 0 h = mitosis; PM indicates post-mitosis). Red box indicates time point shown in (C). (C) Individual z-sections at 1.2 h before mitosis. Arrowheads indicate three fluorescently labelled genomic loci, whereby the filled arrowheads indicate the optical section with highest fluorescence signal. (D-H) Image analysis procedure. (D) 3D maximum intensity z-projection of cell shown in (C). Boxes indicate regions containing labelled alleles. (E) Single z-section containing brightest dCas9-mEGFP signal for the respective alleles. (F) Gaussian filtering of (E). (G) Fluorescence density distribution of (F). (H) Gaussian mixture model fitted to (G). d indicates distance between the means of two Gaussian functions. Scale bars, 10 μm (A and B), 5 μm (C and D) and 1 μm (E).

In G2 nuclei, the fluorescently labelled genomic sites appeared either as a single dot (“singlet”) or as a pair of dots (“doublet”) (Fig. 1, C-E). Singlets represent unreplicated genomic loci or replicated loci where sister DNA strands are at a distance below the resolution limit.

Doublets represent replicated sister loci that are spaced apart beyond the resolution limit. To determine the centre positions of fluorescent dots, we extracted image sub-regions around each singlet or doublet for automated analysis. Owing to the low z-sampling rate, which could not be increased because of phototoxicity limitations, we could not accurately measure distances in 3D space. We therefore performed all distance measurements in 2D optical sections that contained the brightest fluorescent signal for the respective labelled allele (Fig. 1C, filled arrowheads; D and E). We reduced image noise by filtering and then fitted a mixture model of two Gaussian functions to the image sub-regions using an automated optimization algorithm (Fig. 1, F-H; see Methods for details) to obtain the coordinates of two fluorescent point sources – in singlets as well as in doublets.

To estimate the accuracy of the image analysis method, we computationally simulated images based on a pair of fluorescence point sources, the microscope’s point spread function, and detector noise (Fig. S2, G-L). We then determined the distance between simulated fluorescent point sources by fitting the Gaussian mixture model. For simulations with distances larger than 300 nm, the measurements matched ground truth with an accuracy of 12.4 ± 14.5 nm (mean ± SD), but the model fitting did not yield accurate results at smaller distances (Fig. S2, G-I). We hence classified fluorescent dots as doublets when the measured distance was larger than 300 nm, or singlets when the measured distance was equal or smaller than 300 nm. This approach provides an objective, consistent, and accurate means to determine the degree of sister locus separation.

To investigate the organization of sister chromatids during G2, we asked whether sister loci maintain a persistent linkage at any of the labelled genomic sites in our cell line collection. We imaged each of the 16 cell lines as described above. For all 16 genomic sites, we found that a fraction of loci appears as doublets during G2, with sister loci spaced up to ∼1.5 µm apart (Fig. 2, A and B). The distance between sister loci during interphase was often as large as in mitotic chromosomes (Fig. 2 C). Hence, none of the investigated genomic sites displayed constitutive sister locus linkage.

**Figure 2.**
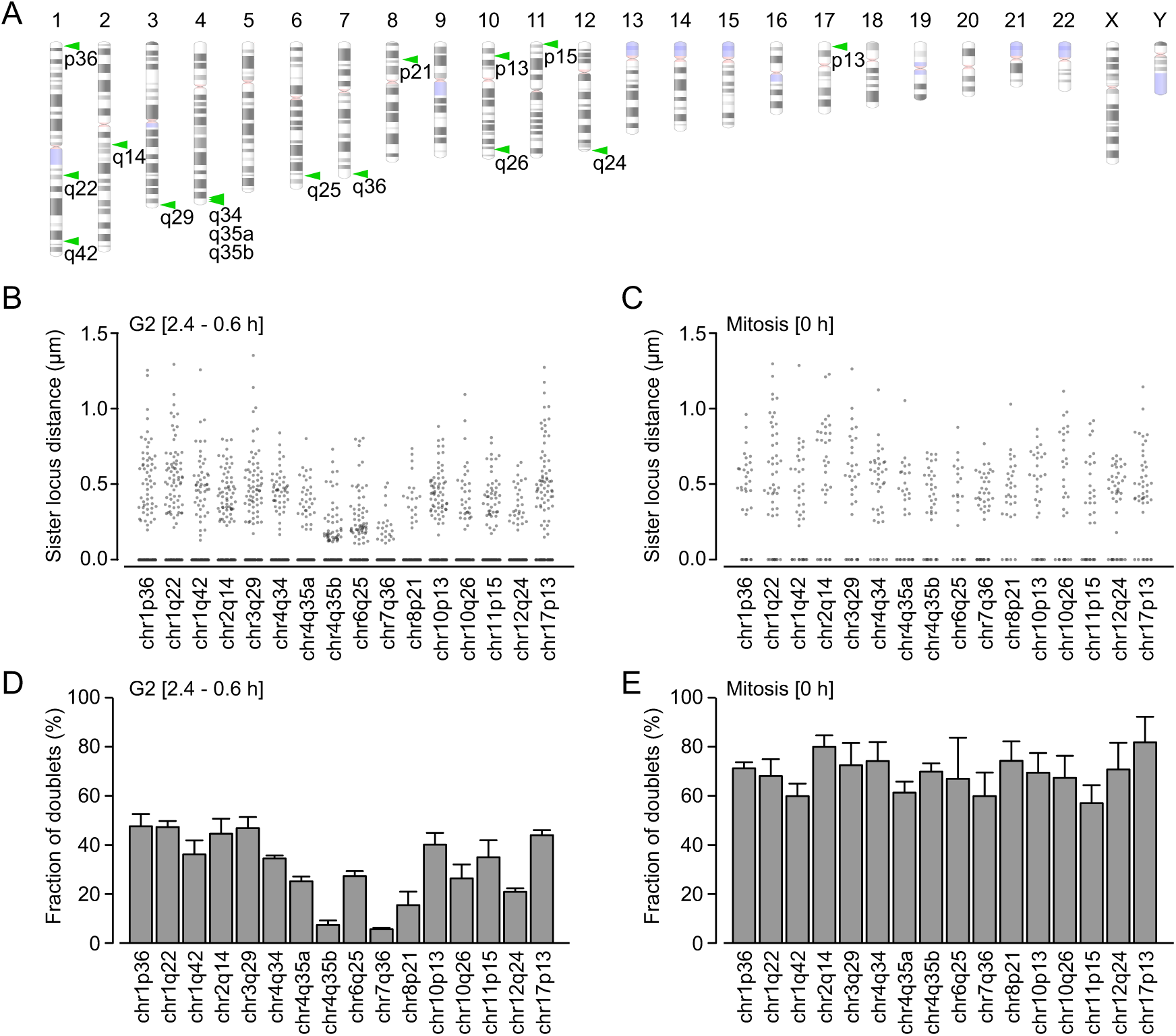
Mapping sister chromatid resolution during interphase in live human cells. (A) Position of dCas9-mEGFP/sgRNA-labelled genomic sites (green arrowheads) contained in the HeLa cell line collection on human chromosome ideogram (Schneider et al., 2017). Annotations indicate chromosome and band numbers. Blue indicates unmapped chromosome regions, pink indicates centromeres. See Table S1 for details. (B-E) Quantification of sister locus separation in 16 cell lines as in (A), based on live-cell microscopy and image analysis as in Fig. 1. (B) Sister locus distance in G2 (0.6 – 2.4 h preceding mitosis). Each dot indicates distance measurement from one allele; n = 120 randomly subsampled distance measurements for each cell line, based on three independent experiments. (C) Sister locus distance as in (B) during mitosis. n ≥ 23 measurements. (D) Fraction of doublets (spots with distance >300 nm) in G2 based on data shown in (B). Significant differences between genomic sites (p < 10-7 by one way ANOVA). (E) Fraction of doublets in mitosis was calculated for the data shown in (C). The fraction of doublets is not significantly different at different genomic positions (p = 0.75 by one way ANOVA). For D and E, bars indicate mean ± SEM; n = 3 experiments.

To investigate whether sister locus separation depends on the genomic context, we calculated the fraction of doublets in G2. This revealed significant differences between genomic sites (p < 10-7 by one-way ANOVA test), ranging from 5.7% to 47.8% doublets (Fig. 2 D). During mitosis, the fraction of doublets was higher, ranging from 57.2% to 82.0% (Fig. 2 E) and it did not differ significantly between different genomic sites (p = 0.75 by one-way ANOVA test). These data suggest that the spatial organization of sister chromatids during G2 depends on local genomic features.

To investigate potential perturbations caused by the in vivo labelling method, we used FISH to visualize six sgRNA target sites in wild-type HeLa cells and in cells expressing dCas9-mEGFP and the respective sgRNA. We synchronized cells to G2 by a release from double thymidine block and performed FISH, which revealed a high correlation of sister locus separation between wild-type cells and labelled cells (Fig. 3, A-D; Pearson correlation coefficient R2 = 0.81). Thus, the dCas9-mEGFP/sgRNA labelling and FISH data support our inference that the topology of sister chromatid separation in interphase depends on the genomic context.

**Figure 3.**
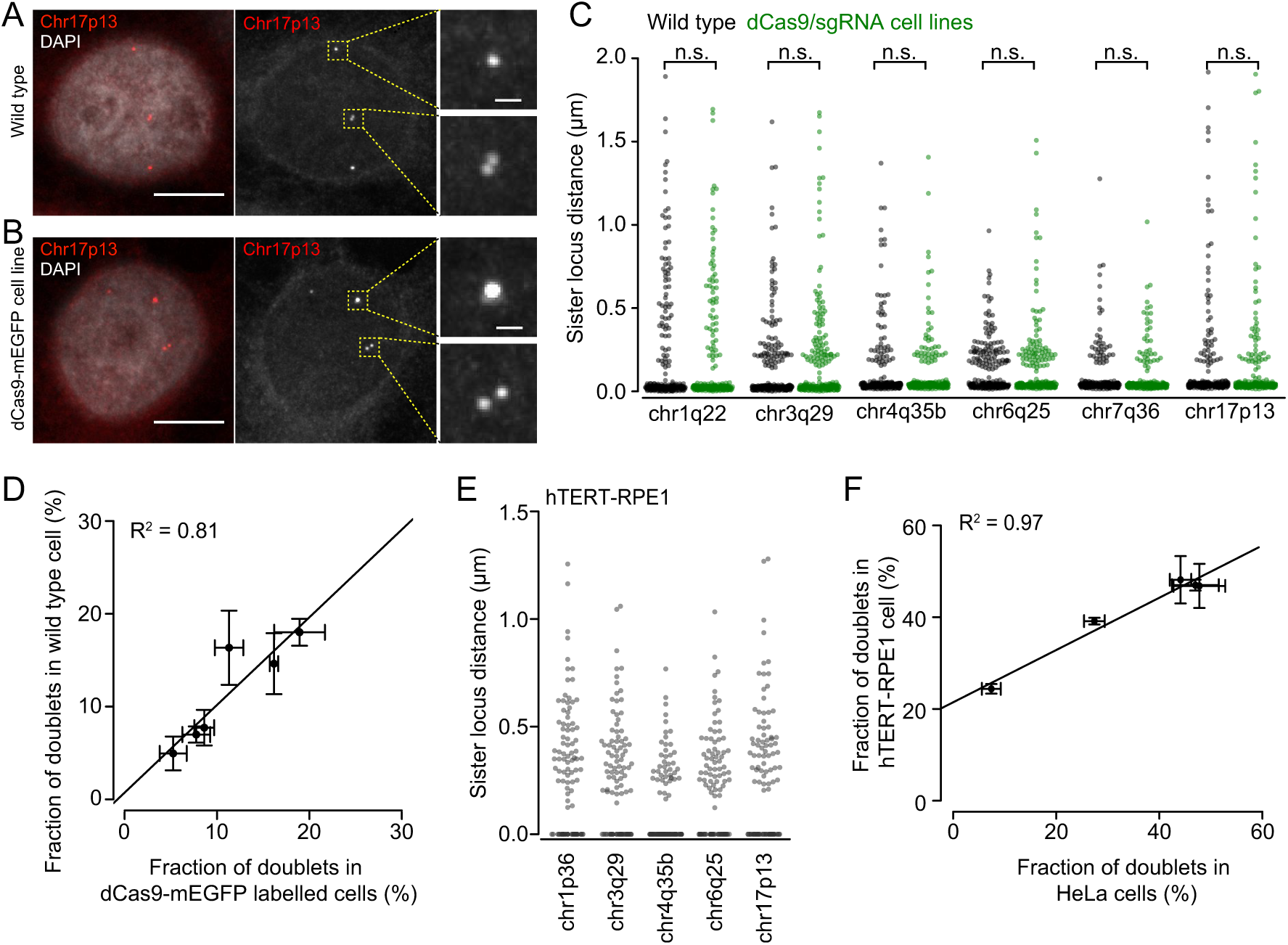
Mapping sister chromatid resolution in G2 in HeLa cells using FISH, and in hTERT-RPE1 cells using dCas9-mEGFP. (A, B) Comparison of wild type HeLa cells and HeLa cells expressing dCas9-mEGFP and sgRNA targeting Chr17p13. Cells were synchronised to G2 by release from double thymidine block and stained with FISH probes targeting Chr17p13. Yellow boxes indicate labelled alleles, as shown in insets. DNA was stained with DAPI. (C) Quantification of sister locus distances based on FISH staining as in (A, B) for six genomic sites, in wild type cells and in cell lines expressing dCas9-mEGFP together with the sgRNA targeting to the corresponding locus. N = 230 loci from 3 independent experiments for each genomic site; n.s. indicates p > 0.05; non-parametric Wilcoxon rank test. (D) Fraction of doublets based on data shown in (C). (E) Sister locus distances in five dCas9-mEGFP/sgRNA labelled hTET-RPE1 cell lines, determined by live-cell microscopy in Fig. 1; n = 100 randomly subsampled distance measurements for each cell line. (F) Fraction of doublets in G2 (2.4 – 0.6 h before mitosis) in hTERT-RPE1 cells based on (E), compared to the fraction of doublets in HeLa Kyoto cells (as in Fig. 2 D). Dots and error bars indicate mean ± SEM; n ≥ 3 experiments. Scale bars, 10 μm; insets: 1 μm.

Finally, we asked whether the locus-specific variation of sister chromatid separation is conserved between different cell types. We generated non-cancer Retinal Pigmented Epithelial (hTERT-RPE1) cell lines expressing dCas9-mEGFP and sgRNA, targeting five different loci (Chr1p36, Chr3q29, Chr4q35b, Chr6q25, Chr17p13, respectively), and imaged them by 3D live-cell confocal microscopy. The extent of sister locus separation in G2 for each chromosomal site was highly correlated between HeLa and hTERT-RPE1 cells (Fig. 3, E and F, Pearson correlation coefficient R2 = 0.97). Thus, the genome specifies the sister chromatid resolution topology in a cell-type-independent manner.

### Proximity to cohesin ChIP-sequencing peaks does not suppress sister locus separation

The degree of sister locus separation might be governed by the distance to genomic cohesin enrichment sites. To investigate this, we calculated the genomic distance between the center point of each dCas9-mEGFP/ sgRNA-labelled locus and the nearest cohesin enrichment site, based on ChIP-sequencing profiles of the SMC3 subunit in G2 HeLa cells (Ladurner et al., 2016) (Fig. 4 A). We then compared this to the degree of sister locus splitting in G2 based on the live cell imaging data shown in Fig. 2 D. Unexpectedly, increased distance from SMC3 ChIP-sequencing peaks did not correlate with an elevated doublet frequency (Fig. 4 B). Instead, loci in genomic proximity to SMC3 ChIP-seq peaks appeared as doublets more frequently than those further away (Pearson correlation coefficient R2 = 0.36), even when very close (the smallest genomic distance of 2.2 kb corresponding to a Euclidian distance of 140 nm for a fully-extended 10 nm chromatin fiber). Hence, sister loci can resolve substantially even when close to genomic cohesin enrichment sites.

**Figure 4.**
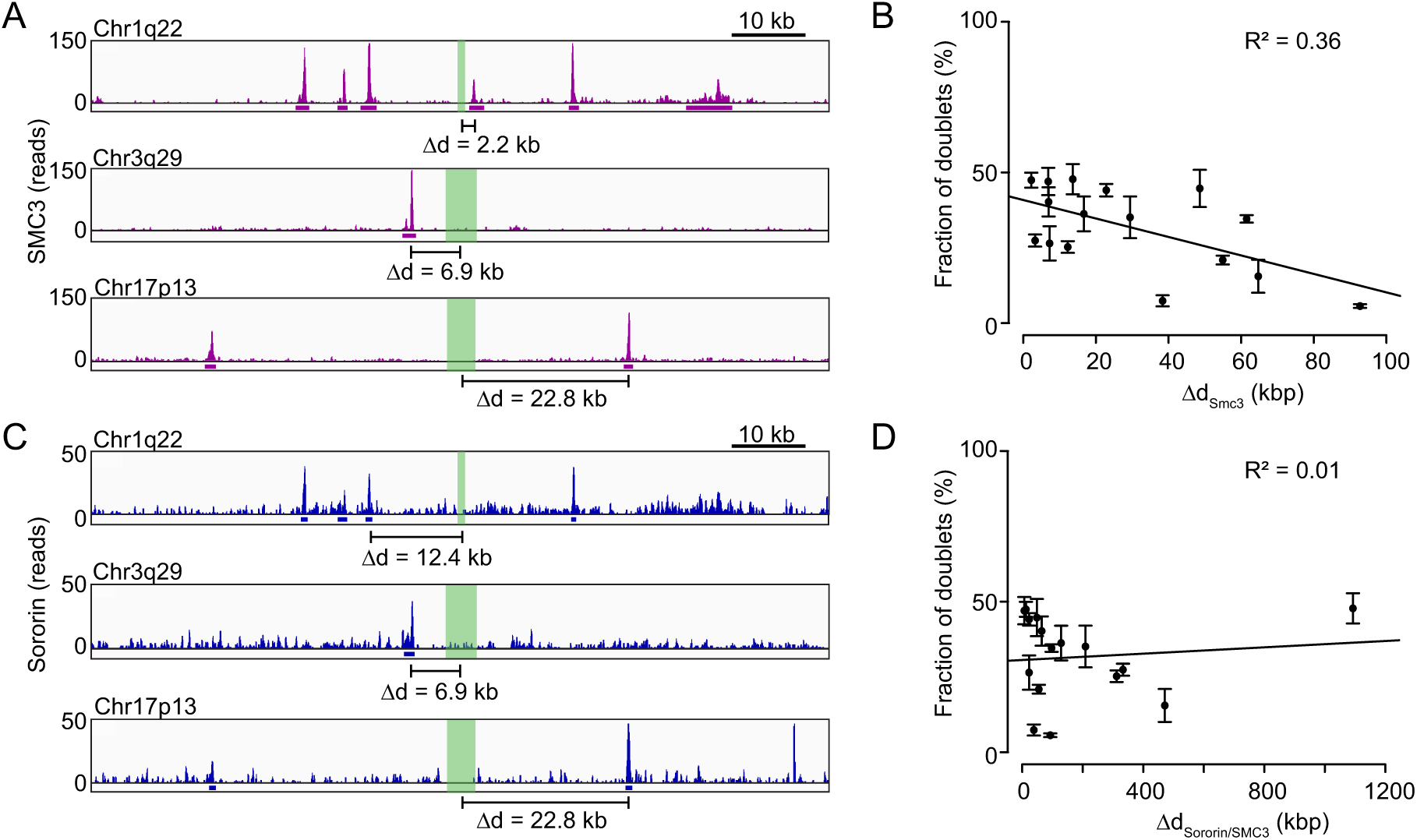
Genomic regions close to cohesin/Sororin ChIP-sequencing peaks frequently separate sister loci during G2. (A) Genomic maps of SMC3 binding profiles around three dCas9-mEGFP/sgRNA-labelled loci. Magenta indicates DNA-sequencing read counts and annotated peaks for SMC3 ChIP-sequencing data from (Ladurner et al., 2016). Green indicates sgRNAs target regions used for dCas9-mEGFP imaging. Genomic distances were calculated between the centre of each sgRNA-target region and the nearest SMC3 ChIP-sequencing peak. (B) Genomic distances between sgRNA-target regions and the nearest SMC3 ChIP-sequencing peak for all 16 genomic sites labelled by dCas9-mEGFP/sgRNA (Table S1), related to the fraction of doublets in G2 as shown in Fig. 2 D. Bars show mean ± SEM; n = 3 experiments. (C) Genomic maps of Sororin binding profiles around the loci shown in (A). Blue indicates DNA-sequencing read counts and annotated peaks for Sororin ChIP-sequencing data from (Ladurner et al., 2016). Green indicates target regions of sgRNAs used for dCas9-mEGFP imaging. Genomic distances were calculated between the centre of each sgRNA-target region and the nearest Sororin ChIP-sequencing peak that also contained SMC3. (D) Genomic distances between sgRNA-target regions and the nearest genomic site containing ChIP-sequencing peaks of SMC3 and Sororin, related to the fraction of doublets in G2 as shown in Fig. 2 D. Bars show mean ± SEM; n = 3 experiments.

We next considered that only a subset of cohesin enrichment sites might contribute to sister chromatid cohesion. Specifically, we asked whether only SMC3 enrichment sites that also contain the cohesin stabilization factor Sororin influence the degree of sister locus splitting. We calculated the genomic distance between dCas9-mEGFP/sgRNA-labelled loci and the nearest SMC3 ChIP-sequencing peak that also contained Sororin in G2 (Ladurner et al., 2016) (Fig. 4 C). This showed that genomic proximity to Sororin/SMC3 ChIP-sequencing peaks also did not correlate with a reduced fraction of doublets (Fig. 4 D). Together, these data suggest that cohesin and Sororin ChIP-sequencing peaks do not represent sites of preferential sister chromatid cohesion – at least at the scale that can be probed by our cell line collection.

### Degree of sister locus separation correlates with nuclear positioning, chromatin state, and DNA replication timing

We next searched for other genomic features that might explain the variations of sister locus separation. To detect potential implications of the chromatin state, we tested whether the degree of sister locus splitting correlates with nuclear positioning, as transcriptionally inactive constitutive heterochromatin localizes to the nuclear periphery, whereas actively transcribed euchromatin predominantly localizes to interior regions of the nucleus (Pueschel et al., 2016). We determined the distances of labelled loci from the nuclear boundary for each of the 16 cell lines (Fig. 5, A-C) and correlated this with the sister locus splitting data (see Fig. 2 D). This showed that genomic sites closely associated with the nuclear periphery generally had a low fraction of doublets in G2 (Fig. 5 D).

**Figure 5.**
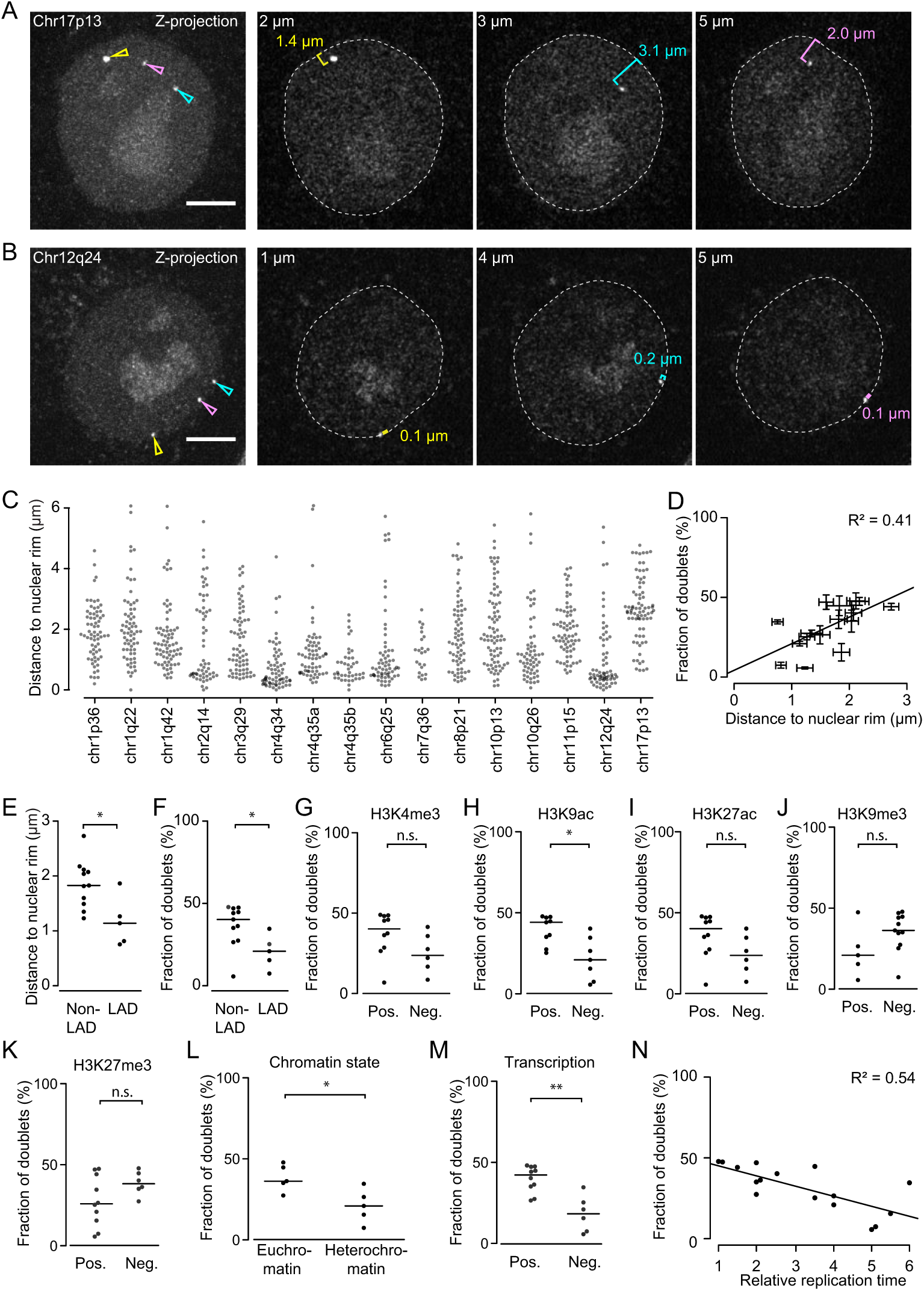
Degree of sister locus separation correlates with chromatin state. (A-D) Nuclear positioning of dCas9-mEGFP-labelled loci. (A) Z-projection and individual z-sections from live cell expressing dCas9-mEGFP and sgRNA targeting Chr17p13. The nuclear rim (dashed line) was determined based on the nucleoplasmic background fluorescence and the distance to dCas9-mEGFP-labelled loci was measured in individual z-sections. (B) As (A) for Chr12q24 locus. (C) Distance to the nuclear rim for 16 genomic loci (Table S1), as in (A, B). Differences between genomic positions are significant (p < 10-16, by Kruskal-Wallis rank sum test; n ≥ 23 measurements per locus). (D) Relationship between locus distance to the nuclear rim and the fraction of doublets during G2 (as in Fig. 2 D). Error bars indicate mean ± SEM; n = 3 experiments for fraction of doublets; n ≥ 23 nuclear rim distance measurements. (E, F) The 16 labelled loci were classified according to their position within or outside of lamina associated domains (LADs) (Guelen et al., 2008) and the respective mean distance to the nuclear rim (E) or fraction of doublets (F) plotted for each locus; * indicates p = 0.02 by unpaired two-sided t test. (G-L) Correlation between sister locus separation and histone modifications in 100 kb regions centred around labelled loci, based on HeLa data from (Hansen et al., 2010; ENCODE Project Consortium, 2012). Pos. indicates the presence of at least one of the respective chromatin marks, Neg. indicates absence. Dots indicate labelled loci. (G-I) Transcriptionally active chromatin marks H3K4me3 (G), H3K9ac (H), and H3K27ac. (J, K) Repressed chromatin marks H3K9me3 (J) and H3K27me3 (K), n.s. indicates p > 0.05 and * indicates p = 0.01 by unpaired two-sided t test. (L) Classification of chromatin state surrounding the labelled loci, based on the presence of only active chromatin marks (H3K4me3, H3K9ac, H3K27ac), or only repressive marks (H3K9me3, H3K27me3), as shown in (G-K); * indicates p = 0.02 by unpaired two-sided t test. (M) Transcriptional activity in 100 kb region centred around labelled loci, based on polyadenylated RNA transcript profiling in HeLa cells (Hansen et al., 2010; ENCODE Project Consortium, 2012), ** indicate p = 0.003 by unpaired two-sided t test. Bars indicate median (E - M). (N) Relationship between sister locus splitting during G2 and DNA replication timing (based on (Hansen et al., 2010), whereby class 1 is earliest and class 6 is latest) for 16 labelled loci.

To further test if sister locus separation correlates with nuclear localization, we considered DNA sequencing-based annotations of genomic regions associated with the nuclear periphery, termed lamina-associated domains (LADs) (Guelen et al., 2008). Of the 16 genomic sites contained in our cell line collection, 5 reside in LADs (Table S1).

These genomic sites indeed localized significantly closer to the nuclear rim in our imaging experiments compared to genomic sites residing outside of LADs (Fig. 5 E). The fluorescently labelled loci residing in LADs had a significantly smaller fraction of doublets in G2 compared to loci residing outside of LADs (Fig. 5 F). Thus, genomic regions associated with the nuclear periphery separate sister loci less frequently than genomic regions residing predominantly in the nuclear interior.

To investigate whether the degree of sister locus separation correlates with chromatin states, we considered HeLa ChIP-seq data of histone modifications (Hansen et al., 2010; ENCODE Project Consortium, 2012). We determined whether 100 kb regions centered around each genomic target site contained marks of transcriptionally active euchromatin (H3K9-acetylation, H3K27-acetylation, and H3K4-trimethylation) or marks of transcriptionally repressed heterochromatin (H3K9-trimethylation and H3K27-trimethylation). We consistently observed that labelled loci that had active euchromatin marks in their vicinity split more frequently, whereas labelled loci that had repressive heterochromatin marks in their vicinity split less frequently (Fig. 5, G-K). For the individual histone modifications these differences were not statistically significant (by unpaired two-sided t-test at α = 0.05), except for H2K9-acetylation, but considering multiple modifications to classify euchromatin (only active marks) or heterochromatin (only repressive marks), we did detect significant differences (Fig. 5 L; p = 0.02 by unpaired two-sided t test). Thus, genomic loci residing in euchromatic regions have a higher probability to separate their sisters than those residing in heterochromatin.

We next investigated whether the degree of sister locus separation correlates with transcriptional activity. We considered mRNA expression profiling data from HeLa cells (Hansen et al., 2010; ENCODE Project Consortium, 2012) and found that sister loci residing in the vicinity of transcribed chromatin were split more frequently than sister loci residing in untranscribed regions (p = 0.003 by unpaired two-sided t test). Hence, open chromatin and transcriptional activity correlate with increased sister locus separation in G2.

Heterochromatin at the nuclear periphery replicates late during S-phase compared to euchromatin of the nuclear interior (Leonhardt et al., 2000). The high incidence of sister locus splitting at genomic sites that preferentially localize to the nuclear interior might hence result from early DNA replication. To investigate this, we related our sister locus separation measurements to published DNA replication timing data from HeLa cells (Hansen et al., 2010; ENCODE Project Consortium, 2012). Our live-cell imaging data of G2 cells (Fig. 2 D) showed that dCas9-mEGFP-labelled loci residing in early-replicating genomic regions appeared as doublets more often than those residing in late-replicating regions (Fig. 5 N). Thus, the spatial organization of sister chromatids correlates with the DNA replication program.

### Dynamics of sister chromatid resolution

We next investigated the dynamics of sister locus separation during cell cycle progression. First, we aimed to determine when sister loci start to split. We followed the trajectories of individual labelled alleles from G1 until mitotic entry using the data shown in Fig. 1. For the early-replicating locus Chr17p13, the first doublets appeared more than 12 h before mitotic entry (Fig. 6, A-D). By 8 h before mitotic entry, doublets had appeared in more than 50% of the trajectories, and by 2 h before mitotic entry in 93% of the trajectories (Fig. 6, C and D; n = 40 trajectories from 27 cells). Thus, sister chromatids at this genomic site can split many hours before cells enter mitosis. Moreover, the time point of initial sister locus separation varies several hours between different alleles and cells.

**Figure 6.**
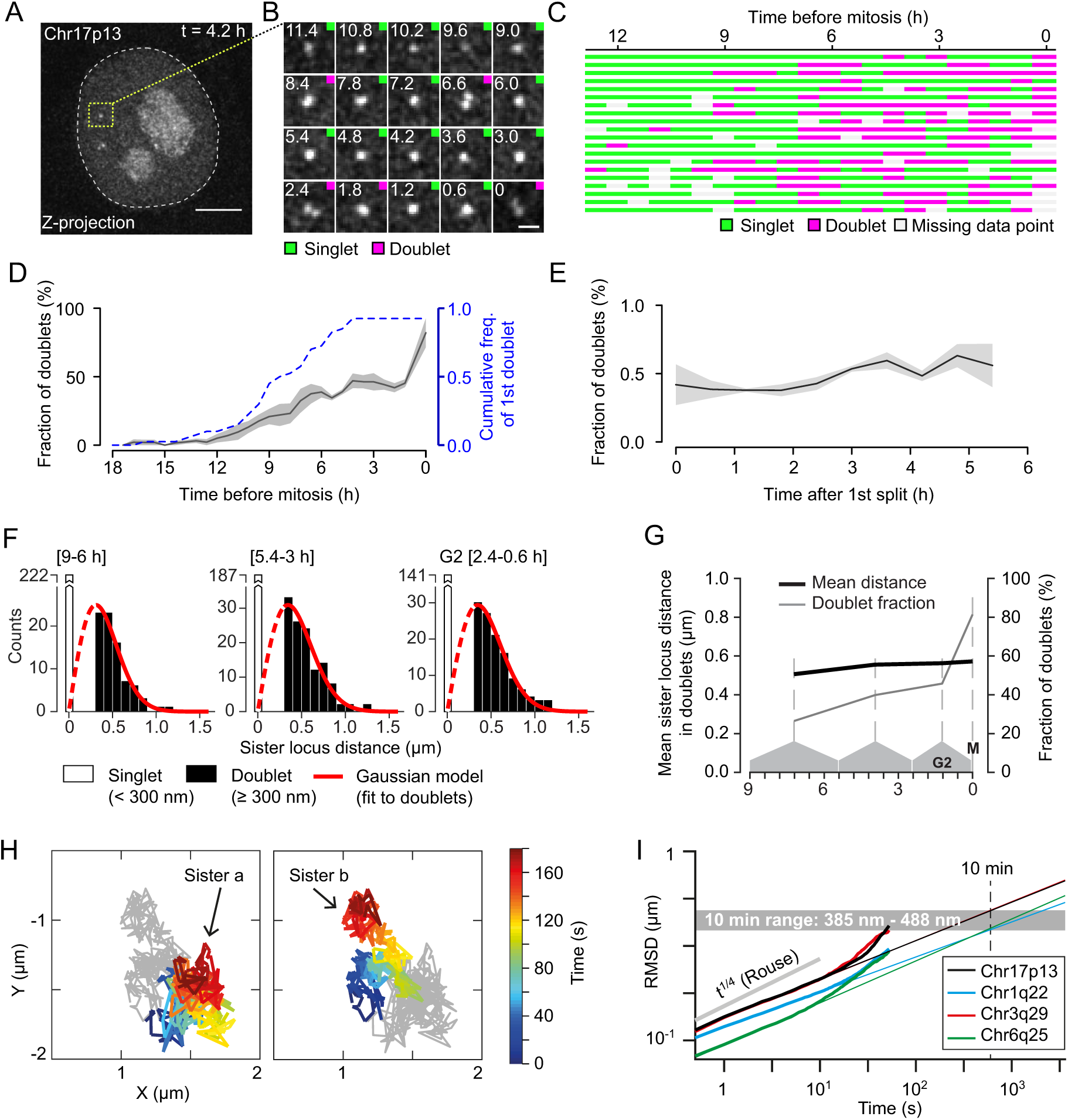
Dynamics of sister locus resolution from S-phase until mitosis for Chr17p13. (A-G) Analysis of individual locus trajectories from imaging data as shown in Fig. 1 and 2. (A) Cell expressing dCas9-mEGFP and sgRNA targeting locus Chr17q13. Dashed white line indicates nuclear rim, yellow box indicates region for trajectory analysis as shown in (B), time points are relative to mitosis (t = 0 h). Coloured squares indicate doublet (magenta) or singlet (green), respectively, based on automated analysis as in Fig. 1 D-H. (C) Twenty trajectories of individual alleles for Chr17q13 as shown in (B). (D) Fraction of doublets based on trajectories as in (C). Solid line and shaded area indicate mean ± SEM; n = 3 experiments. Dashed line shows cumulative frequency of first doublet detected in individual trajectories; n = 40 trajectories. (E) Fraction of doublets as in (D) after aligning all the trajectories to the 1st observed doublet. (F) Distribution of sister locus distances for data shown in (A-E) at indicated cell cycle periods and Rayleigh distributions (corresponding to a Gaussian model) fit to the doublet distance distributions. (G) Mean distance between sister loci in doublets, based on data shown in (F). (H, I) Diffusional mobility of dCas9-SunTag-labelled genomic sites. Cells synchronized to S/G2 were imaged with a time-lapse of 0.5 s and doublets were considered for diffusional mobility analysis. (H) Representative trajectories for Chr17p13. Coloured trajectory indicates one of the sister loci; grey trajectory indicates the respective other sister. (I) Root mean squares displacement (RMSD) analysis for Chr17p13 and three other genomic sites. Thick lines indicate measured data, thin lines indicate extrapolations based on the initial curve segments up to 10 s. Grey line indicates mobility of a Rouse polymer model. Scale bars, 5 μm (A) and 1 μm (B).

After the initial split, sister loci might remain separate until cells enter mitosis. However, analysis of individual allele trajectories showed that doublets frequently alternate with singlets before cells enter mitosis (97 % of the trajectories for Chr17p13; n = 37; see Fig. 6 C). The alternation between singlets and doublets, however, does not imply that sister loci switch between linked and dissociated states, as our imaging set-up resolves fluorescent dots only above distances above ∼300 nm along the x-y image plane. Furthermore, singlets might represent sister loci that separate even further apart along the z-axis, which we cannot reliably quantify owing to the low optical sectioning along the z-axis. Nevertheless, these data show that replicated interphase chromosomes form a dynamic structure in which sister loci continuously move.

While the cell population progressed towards G2, the incidence of doublets gradually increased to about 50%, and upon mitotic entry to around 80% within a single time frame of 36 min (Fig. 6, C and D). This is still below the cumulative frequency of first split measured in single allele trajectories at late interphase stages, consistent with dynamic alternations between singlets and doublets. The gradual increase of doublets during interphase progression might be explained by the variable onset of sister locus splitting in different cells or alleles or by a continuous and slow drift separating sister loci. To investigate this, we re-aligned all trajectories to the time of first doublet appearance. We found that following the initial split, the incidence of doublets did not increase further (Fig. 6 E). This is consistent with a single rapid process releasing sister loci into separated sister chromatids.

To investigate if sister loci continue to move further apart once they have been released, we analyzed distances between separated sister loci in doublets at different periods of interphase. While the fraction of doublets increased as cells progressed towards G2, the mean distances between sister loci in doublets barely changed (Fig. 6, F and G). This is consistent with a model where sister loci separate by a single event and then rapidly equilibrate their distances within dynamic sister chromatids.

To further investigate this, we tested whether the sister locus separation measurements fit to a mathematical model of dynamic equilibrium polymers linked by cohesin complexes (see methods for details). The dynamic equilibrium model predicts that the distribution of relative sister locus positions in 3D nuclear space follows a 3D Gaussian function. In 2D image projections, as in our experimental microscopy data, the distribution of sister locus distances according to this model thus follows a Rayleigh distribution p(d) = d/s2 exp(-d2/2s2), where d is the 2D sister locus distance and the only parameter s determines the scale of the distribution. We fitted Rayleigh functions to the distance distributions of doublets at different cell cycle stages and obtained very good agreement (Fig. 6 F, red lines). The observed sister locus distances are hence consistent with a model where sister loci rapidly equilibrate their relative positions after the initial split.

A dynamic equilibrium model relies on the assumption that sister loci explore their equilibrium distances quickly after replication. In order to assess this, we considered the diffusional mobility of individual genomic loci. To improve signal-to-noise for high time-resolution imaging experiments, we generated a HeLa cell line expressing dCas9 tagged with the SunTag system (Tanenbaum et al., 2014). We individually introduced sgRNAs targeting Chr17p13, Chr1q22, Chr3q29, or Chr6q25 to generate four different cell lines, which yielded ∼4-fold brighter dots compared to the respective dCas9-mEGFP-labelled loci (example shown in Fig. S2, M-O). We synchronized cells to S phase by collecting mitotic cells and growing them for 12 h and recorded confocal microscopy movies at two frames per second. We automatically tracked sister loci and identified trajectory runs containing doublets for at least 10 s length to compute root mean square displacements (RMSDs) for individual sister loci (Fig. 6, H and I). RMSDs increased with time as a power law RMSD = A tα/2 with an exponent α/2 of ∼0.2 and A ∼0.13 µm / sα/2, consistent with previous observations in B lymphocytes (Lucas et al., 2014). To estimate the space explored by loci within 10 min, we extrapolated the power law determined at 0.5-10 s, and found root mean square displacements (RMSD) ranging between 390 and 490 nm for the different genomic loci, comparable to the mean sister distances in doublets. Taken together, these data are consistent with a dynamic chromatin model where after initial separation sister loci quickly equilibrate their relative positions.

We next investigated the dynamics of sister chromatid resolution at other genomic positions. We first studied nine additional cell lines with fluorescently labelled loci that replicate in the first half of S-phase. For all of these loci, we observed a substantial fraction of doublets 12-9 hours before mitotic entry and multiple alternations between singlets and doublets in individual allele trajectories (Fig. 7, A and B; Fig. S3, A-I). The distance between sister loci in doublets did not increase during interphase progression (Fig. 7 C) and the fraction of doublets did not increase after the initial split for any of the genomic sites (Fig. 7, D). Overall, these data indicate that sister loci at various early-replicating genome regions resolve with similar kinetics, albeit to a variable extent.

**Figure 7.**
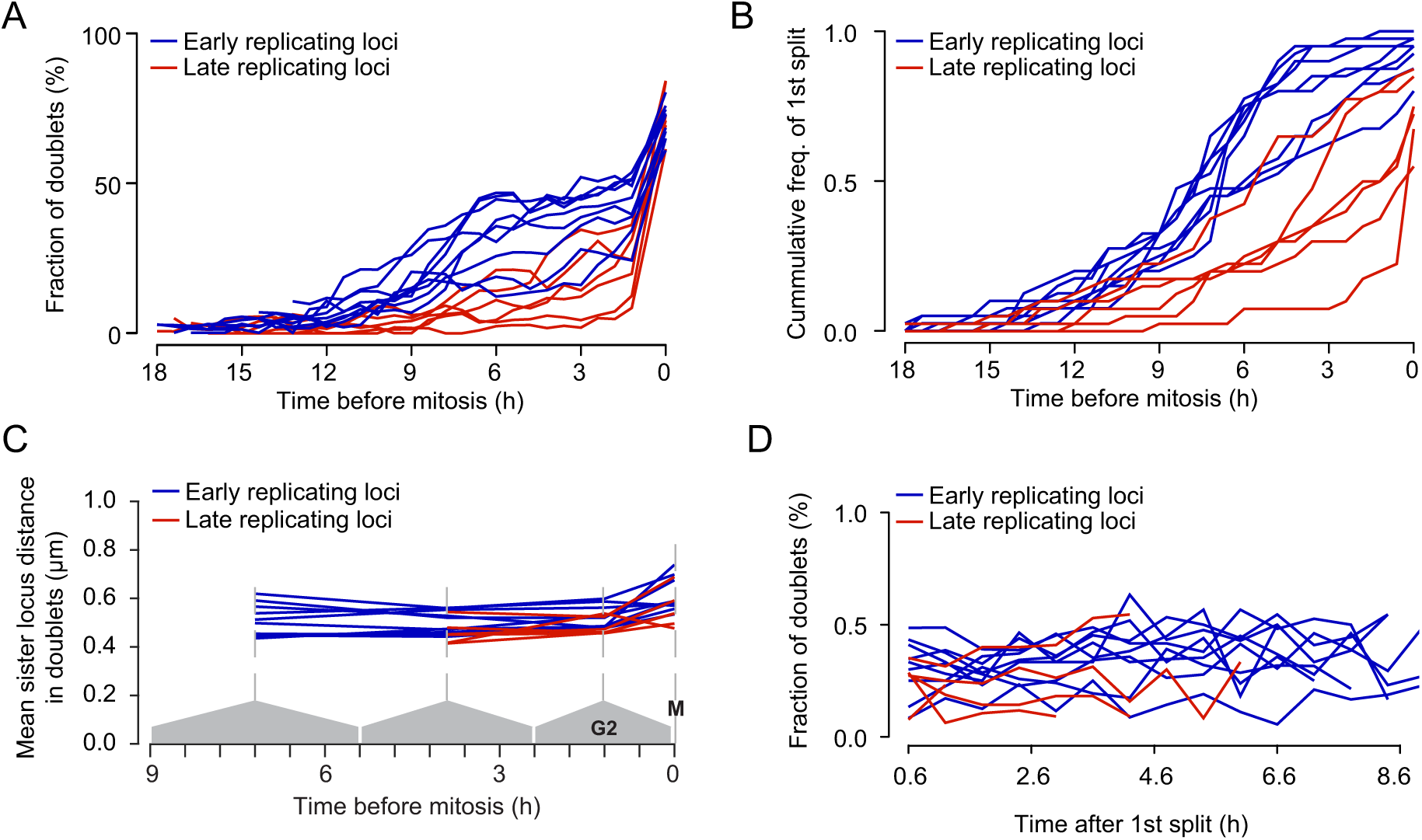
Dynamics of sister locus resolution from S-phase until mitosis for 15 genomic sites. (A) Fraction of doublets as in (Fig. 6 D) for 9 early-replicating (blue) and 6 late-replicating (red) loci. Full data in Fig. S3, A-I. (B) Cumulative frequency of first split event for data shown in (A), see Fig. S3 J-O. (C) Mean distance between sister loci in doublets, calculated as in (Fig. 6 G). (D) Fraction of doublets in trajectories aligned to the 1st doublet for loci shown in (A).

We investigated six cell lines with labelled genomic sites residing in late-replicating chromosome regions and found that the doublet fraction for these loci increased substantially later than observed in early replicating loci (Fig. 7 A; Fig. S3, J-O). However, at 2.4 h before mitosis, most of the individual trajectories had shown at least one doublet for any of the labelled genomic sites (Fig. 7 B; Fig. S3, J-O), confirming that DNA replication was largely complete (with the exception of Chr7q36, see Fig. S3 L). Late-replicating loci did not move further apart and the probability of separation did not increase after their initial split (Fig. 7, C and D). Hence, all investigated genomic sites show consistent patterns of sister chromatid resolution kinetics.

### Sororin delays sister locus separation during interphase

Our observations suggest that parts of sister chromatids move apart during G2 as far as in mitotic chromosomes. This raises the question how cohesin influences the spatial organization of sister chromatids during interphase. To investigate this, we depleted Sororin, which is essential to maintain sister chromatid cohesion (Schmitz et al., 2007; Ladurner et al., 2016). Transfection of siRNA targeting Sororin induced a pronounced mitotic arrest (Fig. S4, A and B), as previously reported (Rankin et al., 2005), validating efficient target protein depletion. The duration of the preceding interphase, however, was not affected by Sororin RNAi (Fig. S4 C), indicating timely DNA replication. The similar duration of interphase enabled us to directly compare locus separation kinetics in Sororin-depleted and unperturbed control cells. By time-lapse microscopy we found that Sororin RNAi caused a premature increase of doublets during interphase for all four investigated genomic sites (Fig. 8, A-F; Fig. S4, D-I), consistent with prior FISH studies (Schmitz et al., 2007; Nishiyama et al., 2010). During late G2, however, the degree of sister locus splitting in control cells approached that of Sororin-depleted cells. Thus, Sororin counteracts sister chromatid separation but it does not mediate permanent linkage of sister loci at any of the sampled genomic sites.

**Figure 8.**
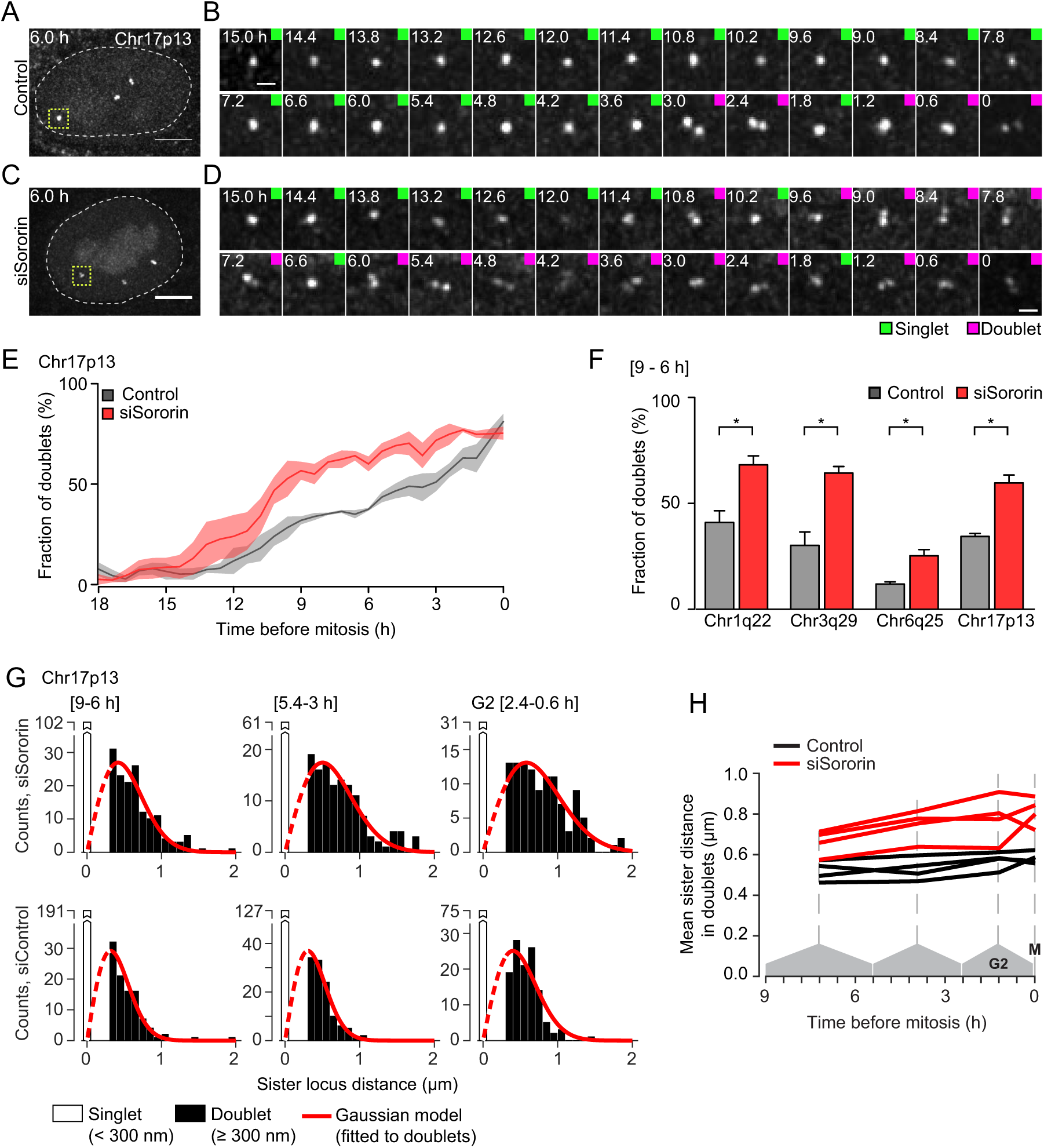
Sister loci resolve faster in Sororin-depleted cells. (A-D) Live cell imaging as in Fig. 1 of Chr17p13 locus labelled by dCas9-mEGFP/sgRNA, after transfection of non-targeting control siRNA (A, B) or Sororin siRNA (C, D). (A) Maximum intensity z-projection of a control cell; dashed line indicates nuclear rim; yellow squares indicates region for (B) trajectory analysis. Time is relative to mitosis (t = 0 h). Doublet (magenta) and singlet (green) were annotated automatically as in Fig. 1 D-H. (C, D) Cell transfected with Sororin siRNA, analyzed as in (A, B). (E) Fraction of doublets over time based on data shown in (A-D). Line and shaded areas indicate mean ± SEM, respectively (n = 3 experiments). (F) Fraction of doublets during S-phase (9 - 6 h prior to mitosis) for data shown in (E) and for three other genomic sites; p = 0.02 for Chr1q22; p = 0.02 for Chr3q29; p = 0.03 for Chr6q25; p = 0.01 for Chr17p13; p values derived from two-sided unpaired t test; n = 3 experiments. (G) Distribution of sister locus distances for data shown in (E) at the indicated cell cycle periods and Rayleigh distributions (corresponding to a Gaussian model) fit to the doublet distance distributions. (H) Mean distance between sister loci in doublets for four genomic sites transfected with siRNA targeting Sororin or non-targeting control siRNA. Scale bars, 5 μm (A, C), 1 μm (B, D).

As for control cells, the distribution of sister locus distances in Sororin-depleted cells fit well to a Rayleigh distribution (Fig. 8 G). The mean distance between sister loci was substantially larger in Sororin-depleted cells compared to control cells at all time points, and it changed only little when cells progressed towards G2 (Fig. 8 H). Hence, premature sister locus separation in Sororin-depleted cells also results in a rapid positional equilibration, whereby the average distances between sister loci are higher than in control cells. This is consistent with a lower abundance of cohesion sites between sister chromatids in Sororin-depleted cells.

## Discussion

Live-cell imaging of endogenous genomic loci has revealed how sister chromatids resolve during progression from S-phase towards mitosis. Our data indicate that the organization of sister chromatids is in part governed by the DNA replication program, as previously suggested (Selig et al., 1992). Sister loci separate early after their replication and rapidly equilibrate their distances in dynamic sister chromatid polymers. In a population of G2 cells, early-replicating loci hence appear split more frequently compared to late-replicating loci. The probability of sister locus separation also correlates with the nuclear localization and chromatin state, consistent with prior observations that transcriptional silencing counteracts sister locus separation (Azuara et al., 2003). Given that chromatin state and nuclear positioning are highly correlated with DNA replication timing, it is difficult to dissect the relative importance of these genomic features for the kinetics of sister locus separation.

The extensive separation of sister loci during S-phase and G2 might result from low abundance of cohesive structures or from potential loop-extrusion activities of cohesin (Fig. 9, A and B). The separation of sister loci might alternatively result from lateral sliding of cohesin along chromatin (Fig. 9 C). In vitro, cohesin can rapidly slide along DNA (Davidson et al., 2016; Stigler et al., 2016) and transcriptional activity can induce relocation of cohesin to distant genomic regions in cells (Lengronne et al., 2004; Busslinger et al., 2017). It is conceivable that cohesin moves from early-towards late-replicating regions of the genome while sister chromatids resolve during interphase progression. The separation of sister loci might further involve a register shift between the two sister chromatids, i.e., with cohesin linking distinct genomic sites in opposing sister chromatids (Fig. 9 D).

**Figure 9.**
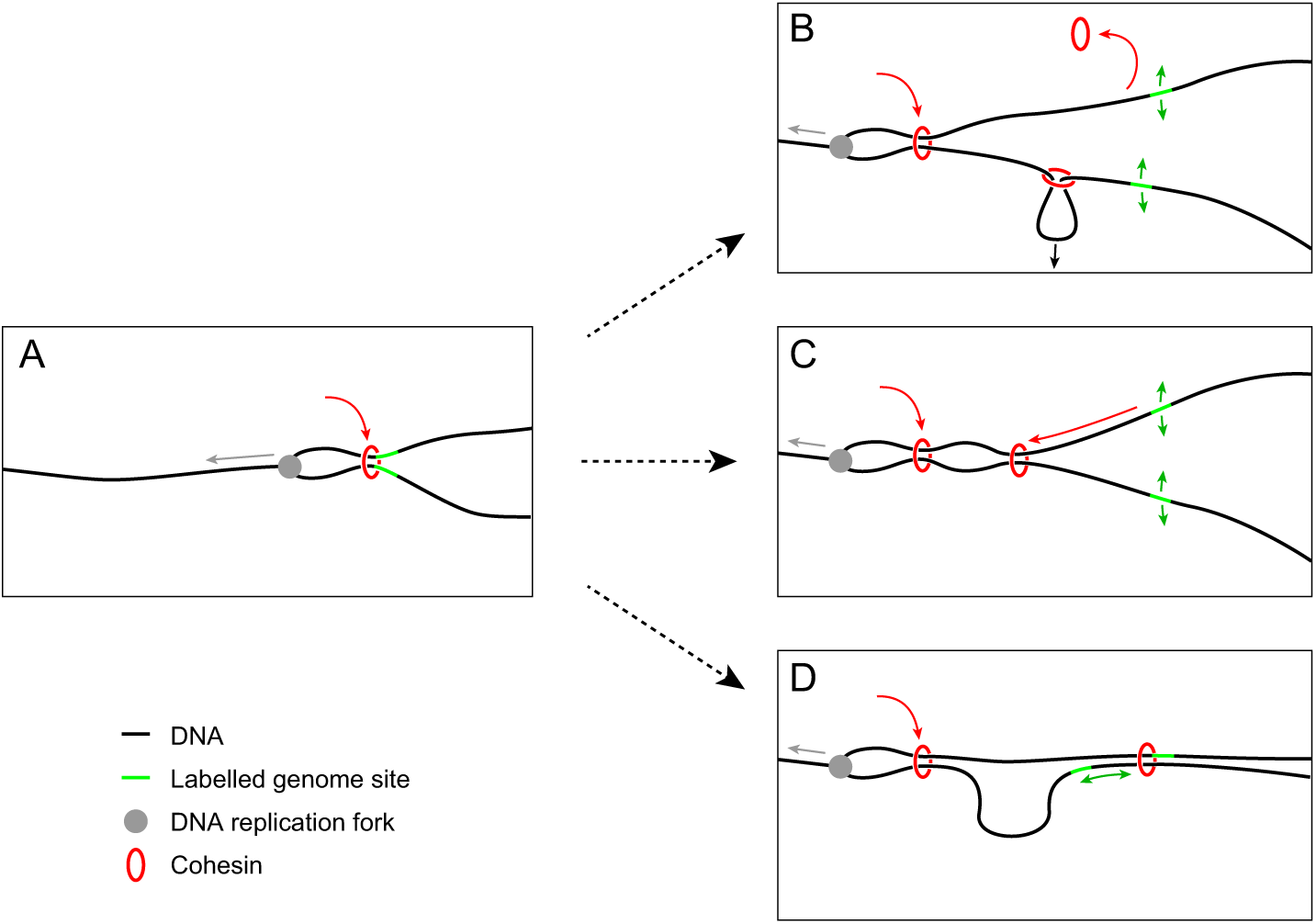
Models for sister locus separation by cohesin reorganization. (A) Sister chromatid cohesion is established co-replicationally. (B) Sister locus separation as a result of cohesin dissociation or cohesin-mediated loop extrusion. (C) Sister locus separation as a result of cohesin sliding along DNA. (D) Sister locus separation as a result of sister chromatid de-alignment.

Overall, our study uncovers how sister chromatids dynamically reorganize during cell cycle progression. It will be interesting to dissect in future studies how chromatin diffusion, topological constraints by cohesion, and potential loop extrusion activities jointly shape sister chromatids. It will also be interesting to investigate the organization of sister chromatids close to centromeres, where cohesion persists for the longest time. Our automated assay system and collection of cell lines with labelled endogenous genomic loci will provide a valuable resource to gain further insights into the mechanics underlying chromosome organization.

## Acknowledgments

The authors thank A. Schleiffer for providing the list of unique repeat sites in the human genome, B. Huang for dCas9-EGFP and gRNA plasmids, H. Zuber, J. Jude, and M. Roth for dCas9-mEGFP HeLa cells, M. Mitter and L. Mirny for comments on the manuscript, IMBA/ IMP/GMI BioOptics and Bioinformatics core facilities for technical support, and Life Science Editors for editing assistance. D.W.G. has received financial support from the European Community’s FP7/2007-2013 under grant agreements no 241548 (MitoSys) and no 258068 (Systems Microscopy), from an ERC Starting Grant (no 281198), from the Wiener Wissenschafts-, Forschungs-und Technologiefonds (WWTF; project nr. LS14-009), and from the Austrian Science Fund (FWF; project nr. SFB F34-06). R.S. has received a PhD fellowship from the Boehringer Ingelheim Fonds. J.N. received financial support from NSF under grant nr. 1504942 (Physics of Chromosomes) and NIH under grant nr. GM114190 (Polymer Models of Mitotic and Interphase Chromosomes). The authors declare no competing financial interests.

## Author contributions

Conceptualization: DWG and RStanyte (lead), JMP (supporting). Data curation: RStocsits. Formal analysis: RStanyte (lead), JN, RH, RStocsits (supporting). Funding acquisition: DWG, RStanyte. Investigation: RStanyte (lead), CB (supporting). Methodology: RStanyte (lead), JN (supporting). Project administration and supervision: DWG. Visualization: RStanyte (lead), JN and RH (supporting). Writing: DWG (lead), RStanyte (supporting).

## Materials and Methods

### Plasmids

For inducible expression of fluorescently tagged catalytically inactive Cas9, the TRE3G-NLS-dCas9-NLS cassette from plasmid pSLQ1658-dCas9-EGFP (Addgene #51023) (Chen et al., 2013) was tagged with mEGFP and sub-cloned into plasmid pQCXIX (Clontech, Cat No.: 631515). For SunTag system, the fluorescently tagged single chain variable fragment that recognises GCN4_peptide (scFV-GCN4-sfGFP) was constitutively expressed from plasmid pHR-scFv-GCN4-sfGFP-GB1-NLS-dWPRE (Addgene #60906) (Tanenbaum et al., 2014). For inducible expression of nuclear localized dCas9 tagged with 10 copies of GCN4 peptide, the NLS-dCas9-2xNLS-10xGCN4_v4 cassette from plasmid pHRdSV40-dCas9-10xGCN4_v4-P2A-BFP (Addgene #60903) was cloned downstream of the TRE3G promoter in the pHR plasmid backbone from Addgene #60906 plasmid. For positive clone selection, the plasmid was further modified to contain the blasticidin S-transferase gene downstream of the cytomegalovirus promoter (pCMV). For efficient sgRNA expression, the sgRNA tracer sequence optimised for imaging (Chen et al., 2013) was cloned into the LentiCRISPR V1 plasmid (Addgene #49535) (Shalem et al., 2014) downstream of the U6 promoter. Additionally, Cas9 and puromycin-N acetyltransferase coding sequences from this plasmid were removed and replaced with the sequence coding for rodent thymocyte differentiation antigen Thy1.1 and neomycin phosphotransferase II fusion protein providing an easy positive clone selection procedure either by FACS or by eukaryotic resistance against Geneticin.

### sgRNA design

The human Tandem Repeat Database (Gelfand et al., 2007) was bioinformatically screened for unique clusters of at least 20 repeats of an at least 20 nucleotide-long sequence that contains a protospacer adjacent motif essential for dCas9/sgRNA labelling (with a minimum conservation between repeats of 80%). Only sequences localizing at a minimum distance of 0.5 Mbp from centromeres and telomeres were considered. SgRNA spacers were designed around available protospacer adjacent motifs avoiding stretches of the same nucleotide and cloned into the sgRNA expression plasmid via BbsI cloning sites, as previously described (Cong et al., 2013).

### Cell lines and Cell culture

Wild type human HeLa cells (‘Kyoto’ strain) were obtained from S. Narumiya (Kyoto University, Japan) and validated by a Multiplex human cell line authentication test (MCA). Human hTERT-RPE1 cell line (further referred as RPE1) was obtained from American Type Culture Collection (ATCC). All cells were cultured in in-house made Dulbecco’s Modified Eagle’s Medium (DMEM), supplemented with 2 mmol L-Glutamine (Gibco), 10% (v/v) fetal bovine serum (FBS) (Gibco) and 1% (v/v) penicillin-streptomycin (Sigma-Aldrich) and grown in a humidified growth chamber at 37°C and 5% CO2. All cells lines in this study were regularly tested for mycoplasma contamination, with negative results.

The parental dCas9-mEGFP and dCas9-10xSunTAG-expressing cell lines were derived from human HeLa Kyoto RIEP and RPE1 RIEP cell line (Samwer et al., 2017) that are modified with rodent-restricted murine ecotropic envelope and allow working with lentivirus at biosafety level 1. Additionally, RIEP cells express Tet3G trans-activator protein (Clontech) for doxycycline inducible gene expression. Cell lines derived from HeLa RIEP and RPE1 RIEP cells were maintained in DMEM medium with 0.5 µg/ml and 5 µg/ml puromycin (Calbiochem), respectively. Parental dCas9-mEGFP expressing HeLa Kyoto and RPE1 cells were generated using lentivirus-mediated DNA delivery as in (Samwer et al., 2017) of the pQCXIX-TRE3G-NLS-dCas9-NLS-mEGFP plasmid packaged in viral particles with ecotropic envelope receptor (EcoR). After viral delivery, a monoclonal dCas9-mEGFP-expressing HeLa cell line was grown from a single colony. The parental dCas9-10xSunTag expressing HeLa Kyoto cell line was generated using the same lentiviral delivery protocol (Samwer et al., 2017) simultaneously infecting cells with viral particles packaged with pHR-scFv-GCN4-sfGFP-GB1-NLS-dWPRE and pHR-TRE3G-NLS-dCas9-2xNLS-10xGCN4_v4-pCMV-BlastR plasmids. Polyclonal parental cell line was derived by selecting with 2 µg/ml blasticidin S (Sigma-Aldrich). All sgRNA-expressing cell lines were further derived from the parental dCas9-mEGFP or dCas9-10xSunTag-expressing HeLa and RPE1 cell lines using the same lentiviral delivery protocol (Samwer et al., 2017). Polyclonal cell lines stably expressing locus-specific sgRNA were selected with 1 mg/mL Geneticin (Gibco). For uniform mEGFP expression levels in HeLa cells, dCas9-mEGFP expression was induced for 48 h with 1 µg/ml doxycycline (Sigma-Aldrich) and low-level-expressing cells were selected using fluorescence-activated cell sorting (FACS). For all experiments dCas9 expression was induced for at least 24 h with 1 µg/ml doxycycline in HeLa Kyoto cells and 1 ng/ml doxycycline in RPE1 cells.

### Cell synchronization

For mitotic shake off, dCas9-mEGFP expression in HeLa cells was induced for 48 h with 1 µg/ml doxycycline (Sigma-Aldrich). 2 h prior to mitotic shake-off, dead and arrested cells were removed by mechanical shake-off and washing cells twice with pre-warmed and CO2-equilibrated PBS. For the following 2 h, cells were grown in pre-warmed and CO2-equilibrated imaging medium (DMEM supplemented with 2 mmol L-Glutamine (Gibco), 10% FBS and 1% Penicillin-streptomycin, but without phenol red and riboflavin). Then, mitotic cells were detached by gently hitting the flask a few times and transferred with the supernatant medium into imaging plates. For fast time-lapse movies cells were imaged 12 – 14 h after mitotic shake off, corresponding to S phase.

For FISH experiments, cells were synchronized to G2 by a double thymidine block. First, cells were arrested at the G1/S transition with 2 mM thymidine (Sigma-Aldrich) for 16 h and released into the cell cycle by washing twice with 37°C and CO2-equilibrated PBS. For the next 8 h, cells were grown in normal DMEM medium as described above. Cells were then arrested for the second time at the G1/S phase transition by 2 mM thymidine for another 16 h and released into the cell cycle by washing twice with PBS, followed by transfer into DMEM. 6 h after the second release from the thymidine block, cells were in G2 phase, as validated by FACS. Throughout the procedure, the medium contained 1 µg/ml doxycycline to maintain dCas9-mEGFP expression.

### Fluorescence activated cell sorting (FACS)

FACS was used to derive live cells with uniform mEGFP expression. Cells were sorted in PBS supplemented with 2% (v/v) fetal bovine serum (FBS) (Gibco), and 2mM Ethylenediaminetetraacetic acid (EDTA) (Applichem) at a FACSAria III (DB) flow cytometer. To assess cell synchronization for FISH experiments, DNA content was measured in cells detached from the plate surface using 0.25% trypsin (Thermo Fisher Scientific), washed twice with PBS and fixed with 73 % ice-cold methanol (Sigma-Aldrich) at -20°C overnight. The next day cells were washed with PBS and stained for 30 minutes at 37°C in a PI-buffer containing 50 ug/ml propidium iodide (Sigma-Aldrich), 10 mM Tris (pH 7.5) (Sigma-Aldrich), 5 mM MgCl2 (Sigma-Aldrich) and 200 µg RNase A (Thermo Fisher Scientific). Cell cycle profiles were analysed with a FACScanto flow cytometer (BD).

### siRNA transfection

RNAi-mediated gene silencing was performed by transfecting siRNA using Lipofectamine RNAiMax (Invitrogen) following manufacturer’s recommendations at room temperature and at a final siRNA concentration of 30 nM. Hs_siSororin (5’ - GCCTAGGTGTCCTTGAGCT-3’) (Schmitz et al., 2007) was used to deplete Sororin and siXWneg9 (5’-TACGACCGGTCTATCGTAG-3’) (Boni et al. 2015) was used as non-targeting control siRNA. For long-term imaging experiments, 8*105 cells were seeded in 75 cm2 flasks and induced with 1ug/mL doxycycline 24-48 h before imaging. Then, cells were synchronised by mitotic shake-off as described above. Mitotic and early G1 cells, where Sororin is degraded (Rankin et al., 2005; Nishiyama et al., 2010), were further transfected with siRNAs and imaged for the next 24 h. In some experiments, unsynchronized cells were imaged (2/3 experiments for chr1q23, 1/3 experiments for Chr3q29 and 1/3 experiments for Chr6q29) 16 h after siRNA transfection.

### Fluorescent in situ hybridization (FISH)

Oligo paint probes (Beliveau et al., 2012) were designed to contain a locus-specific sgRNA sequence at their 3’ end and to have 72°C melting temperature (Table S1). To enhance the fluorescent signal, the sequence at the 5’ end was hybridized with a secondary universal probe (5’-CACACGCTCTTCCGTTCTATGCGACGTCGGTGtttttttt-3’). The probes were synthesized at Integrated DNA technologies (IDT) with a 5’ conjugated fluorophore (ATTO647N) and purified using HPLC. The secondary probe had conjugates with ATTO647N fluorophores on both sides.

For FISH experiments, cells were grown on SuperfrostTM Ultra Plus slides (Thermo Fisher Scientific) and synchronized to G2 using a double thymidine block (see above). Cells were fixed with 4% (v/v) formaldehyde (Thermo Fisher Scientific) for 10 min, washed with PBS, then with double-concentrated saline-sodium citrate buffer 2xSSCT (0.3 M NaCl buffer, 0.03 M trisodium citrate, 1% (v/v) Tween-20) for 5 minutes, then with 2xSSCT buffer containing 50 % (v/ v) formamide (Promega). DNA was denatured in 2xSCCT + 50% (v/v) formamide buffer at 92°C for 2 min and cells were washed for another 20 minutes at 60°C in the same buffer. Cells were covered with hybriwells (Grace Bio-labs) and 1.5 µM of both primary and secondary probes were added in 2xSSCT buffer supplemented with 50% (v/v) formamide, 10% (w/v) dextran sulfate (Sigma-Aldrich) and 20 ng/ml RNase A (Thermo Fisher Scientific). The probes were hybridised in a humidified chamber at 42°C overnight. The next day the slides were washed in 2xSSCT buffer at 60°C for 15 minutes, then in the same buffer at room temperature for 10 minutes and desalted in 0.2xSSC buffer at room temperature for another 10 minutes. Cell nuclei were stained with 1 µg/mL DAPI (Thermo Fisher Scientific) in 0.2xSSC buffer for 1 h at room temperature. Slides were mounted with the anti-fade mounting medium Vectashield (Vector) and stored at 4°C until imaging.

### Microscopy

For all experiments, except of the data shown in Fig. S2, A-F and Fig. 6, H-I, cells were imaged with a scanning confocal Zeiss LSM 780 microscope equipped with a highly sensitive GaAsP detector (Zeiss), and a 63x, 1.4 N.A. Oil Plan-apochromat objective (Zeiss). Cell nuclei were imaged in 11 µm sections with 0.5 µm z spacing and 101 nm pixel size. For whole cell cycle movies, the microscope’s ZEN 2011 software was additionally controlled using an open source macro AutofocusScreen version 3, developed in Jan Ellenberg’s group (European Molecular Biology Laboratory (EMBL), Heidelberg, Germany), which automatically directed the microscope to image the same selected locations every 36 min for 24 h and maintained the focus throughout the experiment. Live cells were imaged in chambered coverslips, using glass bottom LabTek II (Thermo Fisher Scientific) or a plastic bottom ibiTreat (Ibidi). Cells were cultured in imaging medium (DMEM medium supplemented with 10% (v/v) FCS and 1 % (v/v) penicillin/streptomycin, but without phenol red and riboflavin to reduce background autofluorescence (Schmitz et al., 2010), supplemented with 1 µg/ml doxycycline.

Cell proliferation and mitotic progression (Fig. S2, A-F) was scored in movies acquired with a wide field Molecular Devices ImageXpressMicro XL screening microscope using a 10x, 0.5 N.A. S Fluor dry objective (Nikon). Cells were grown in µCLEAR plastic bottom black 96-well imaging plates (Greiner). All cells were treated with 1 µg/ml doxycycline for 48 h before the start of the imaging. 2 h before the start of the imaging, cell nuclei were stained with 125 nM SiR-Hoechst DNA dye (Lukinavičius et al., 2015). To assess mitotic duration, cells were imaged every 4.5 min for 24 h. To score cell proliferation, cells were imaged only 2 times with 24 h interval.

Fast time-lapse movies for diffusional mobility measurements (Fig. 6, H-I) were acquired with a spinning disc confocal UltraView Vox Axio observer (PerkinElmer), using a 150x, 1.35 N.A. glycerol objective (Zeiss) and a Hamamatsu EMCCD 9100-13 camera. The microscope was controlled using Velocity software. Loci were imaged in 7 z-slice sections spaced 0.5 µm for 5 minutes every 0.5 seconds. During imaging, live cells were maintained at 37°C temperature in a humidified 5% CO2 atmosphere.

### Single allele tracking over the cell cycle

Microscopy images were processed using the open-source software Fiji (version 1.48h) (Schindelin et al., 2012). dCas9-mEGFP-labelled genomic loci were tracked throughout the cell cycle using a semi-automated Fiji macro. The first frame of a mitotic cell was manually identified by cell rounding and dispersion of nuclear dCas9-mEGFP signal after nuclear envelope breakdown. Then, two randomly selected individual alleles from each cell were tracked from the beginning of the movie until the following mitosis. Only alleles that were at least 3 µm away from other labelled alleles were considered. 30 x 30 pixel images were cropped and saved for further processing from a single z plane rather than the maximum intensity projection of all z-sections. In case a labelled locus was visible in several z-sections, the slice where two sister loci were visible was selected; otherwise, the z slice with the brightest fluorescence signal was selected. If two fluorescent dots were visible in separate z-sections and it was not possible to distinguish, whether they are two individual sister loci or a single fluorescent dot recorded twice due to its motion, these frames were omitted from the analysis to avoid false-positive singlet detection.

### Quantification of inter-chromatid distances and the fraction of split

Sister chromatid distance was measured using a custom-developed R script based on packages “seqinr” (Charif and Lobry 2007), “spatstat” (Spatial point patterns: methodology and applications with R, 2015), and “SDMTools” (created by Jeremy VanDerWal). Original images were converted into text images using Fiji. The shot noise was then reduced using 0.8 pixel diameter Gaussian blur and further removed by subtracting 95% of the lowest pixel values. The inter-chromatid distance in the images was measured by fitting a mixture of two two-dimensional (2D) Gaussian functions, where x1, y1 and x2, y2 are the centre positions of two sister loci respectively, z1 and z2 amplitudes of the two Gaussian functions approximating the fluorescence intensities of two fluorescent foci and σ2 is the variance of a single Gaussian function, an approximation for the diameter of the microscope’s point spread function. It was assumed, that two sister loci are present in doublet as well as singlet images and therefore a mixture to two 2D Gaussian functions was fitted to all the images. The differences in signal intensity between sister loci was limited to three-fold (z2 ≥ 0.33z1). Additionally, it was assumed, that both sister loci are of the same diameter and are symmetrical in x and y directions (σ = σx1 = σy1 = σx2 = σy2). The 2D Gaussian variance σ2 was measured experimentally, by fitting a single 2D Gaussian function to images of 100 nm diameter fluorescent beads, acquired with the same microscopy settings. To initialize the fitting procedure, approximate centre positions of the fluorescent foci were calculated as the coordinates of the two highest local maxima. It was visually monitored, if both detected local maxima were inside fluorescent foci. If a single bright noise pixel was detected as a maximum, no measurements were recorded. For this, the cell cycle stage was blinded. A mixture of two 2D Gaussian functions was optimized using least square regression analysis.

### Microscopy image simulations to validate spot detection method

To test the accuracy of the image analysis method for quantification of sister locus distances, we simulated microscopy images with two fluorescent dots (related to Fig. S3) using the Python language. First, the positions of the simulated sister loci (µx and µy) were randomly generated within the constrained circular area of 0.3, 0.5 or 0.75 µm diameter. The point spread function of the microscope was approximated with a 2D Gaussian function Mean amplitude A (intensity) ± s.d. and mean variance σ2 ± s.d. for the 2D Gaussian function were calculated from fits of actual microscopy images of Chr3q29 locus. The noise in the images was simulated by inverse transform sampling i.e. drawing a random value for each pixel from the distribution of the noise intensities in actual microscopy images of Chr3q29 locus. Simulated images were further used to determine the centre positions of the two fluorescent dots using the mixture of two 2D Gaussian functions (see above).

### Quantification of locus distance to the genomic cohesin or Sororin enrichment site

Genomic positions of SMC3 and Sororin enrichments sites were calculated for ChIP-sequencing data from HeLa Kyoto cells synchronized to G2 (Ladurner et al., 2016). Peaks were filtered with a p-value threshold of 1e-10 after peak calling with MACS 1.4.2 algorithm, and annotated if present in two independent experimental replicates. Genomic distances were calculated between the centre of each fluorescently labelled locus (centre position of sgRNA target region) and the centre of the nearest SMC3 or SMC3/Sororin ChIPseq peak, respectively, based on GRCh37/hg19 genome assembly.

### Quantification of nuclear positioning for dCas9-mEGFP-labelled loci

The distance of dCas9-mEGFP-labelled loci from the nuclear periphery was measured during S-phase (15 h after mitosis), using confocal long-term imaging data of the entire cell cycle. The distance of dCas9-mEGFP-labelled loci and the nuclear rim (based on nucleoplasmic background fluorescence) was manually measured using Fiji point tool in a single z sections. All visible alleles were measured in a given cell. Two bottom and two top z-sections (1.4 µm each) of the nucleus were omitted as the nuclear edge could not be accurately detected along the z-axis owing to the low z-sampling rate. For doublets, the centre position was used to calculate the distance to the nuclear rim.

### Quantification of histone modification and transcriptional activity

Histone modification data for HeLa cells were obtained from (Hansen et al., 2010; ENCODE Project Consortium, 2012). Gene expression omnibus (GEO) sample accession numbers for individual modifications are as follows: H3K9me3 (GSM1003480), H3K27me3 (GSM733696), H3K4me3 (GSM733682), H3K9ac (GSM733756) and H3K27ac (GSM733684). ±50 kb regions from the centres of the labelled repeats were analysed for the presence of relevant histone modifications. The region was considered positive for the modification if a single peak with p-value smaller than the threshold of 1e-11 was present.

Transcriptional activity in the same ± 50 kb regions around the centres of the labelled repeats was analyzed from polyadenylated RNA sequencing dataset in HeLa cells (Hansen et al., 2010; ENCODE Project Consortium, 2012) with GEO sample number GSM958735. The region was considered positive for transcription if at least one locus in the region was detected in two experimental replicates with a density of 2 reads per million.

### Quantification of labelled allele numbers in each cell line

The number of the fluorescent foci was counted in live-cell 3D confocal images extracted from long term movies (12-18 h after mitosis, corresponding to S - G2 phase). Cell nuclei were observed in multiple time frames and the number of fluorescent nucleoplasmic foci that are visible at least in 3 time points were counted. If two fluorescent foci were less than 2 µm apart and changed from a doublet to a singlet appearance over time, they were considered as sister loci of the same replicated chromosome and counted as a single allele.

### Quantification of cell proliferation and mitotic timing

Cell proliferation was assessed as a fold change in cell number in 24 h. Cell numbers were manually counted using the ‘cell counter’ plugin in Fiji (ImageJ version 1.48v). Mitotic duration was scored manually counting the number of timeframes with chromatin morphologies specific to mitosis (from prophase to anaphase).

### Quantification of signal intensity at dCas9-mEGFP and dCas9-SunTag labelled loci

Stable cell lines with dCas9-mEGFP and dCas9-SunTag labelling Chr17p13 locus were imaged using identical microscopy conditions. Mean fluorescence intensity of a 6×6 pixel region of interest around a labelled locus was measured as signal. An average fluorescence intensity of five mean fluorescence intensity measurements at randomly selected locations in the nucleus using same size region of interest was measured as nucleoplasmic background and subtracted from the signal.

### Dynamic equilibrium polymer model for sister chromosomes

To model sister chromosomes after replication, we assumed two polymers that are linked by cohesin at specific linkage points along the genome. The section between two linkage points is thus a polymer ring. The two sister loci labelled in our experimental tracking data correspond to two specific positions on this ring.

In thermodynamic equilibrium, the distribution of relative positions of two monomers in an ideal polymer ring is a three-dimensional (3D) Gaussian (Khokhlov and Grosberg, 1994). In a 2D projection, like our microscopy data, the distribution of relative positions is a 2D Gaussian, p(r) = 1/(2 π s2) exp(-r2 / 2s2), where r is 2D vector connecting sister loci positions and s is the scale of the distribution. Integrating out the angular orientation, one obtains for the distribution of relative distances d = |r| the Rayleigh distribution p(d) = d/s2 exp(- d2 / 2s2).

Our assumption of thermodynamic equilibrium is based on our findings that (i) neither the fraction of doublets after the first doublet occurrence, nor (ii) the mean sister distance in doublets increases significantly over time, consistent with a picture where sister loci equilibrate after replication on a timescale that is fast compared to cell cycle times, and also consistent with the analysis of mean square displacements. A posteriori, our assumption is validated by a good agreement between our equilibrium model and the data.

Our assumption of ideal polymer rings relies on the Flory theorem in polymer physics, which states that excluded volume effects can be neglected in a dense melt of polymers, which is a reasonable approximation for chromatin in the cell nucleus.

### Histograms of sister distances and fit by Rayleigh distributions

Measured sister distances across several time points were accumulated as indicated by the cell cycle periods in Fig. 6 G. We then binned these data in 100 nm bins. All distances below 300 nm were classified as singlets owing to limited microscopy resolution (see Fig. S2, G-L). We fitted the data with Rayleigh distributions with an additional numerical prefactor A, namely p(d,s,A) = A d / s2 exp(-d2 / 2s2). We introduced the prefactor A instead of normalizing the distribution of distances to unity to preserve the information on sample numbers in the histogram plots. For fitting, we used only doublets (bins with d >300 nm), centred at the respective bin midpoints (350 nm, 450 nm, …). We then minimized the sum of squared differences between p(d,s,A) and the binned data over the parameters A and s. Mean sister locus distances in Fig. 6 G and J are computed only for doublets (data with d >300 nm).

### Mean square displacements and diffusional mobility analysis

Root mean square displacements (RMSDs) were computed from tracking data of movies recorded with two frames per second. The 2D positions of sister loci were automatically determined in maximum intensity projection images by fitting mixtures of two Gaussian functions (see above). Individual sister loci were tracked using nearest neighbour tracking by minimizing the sum of both sister loci displacements in two consecutive frames. Diffusional mobility analysis was then performed on consecutive frames where sister loci appeared as doublets. Trajectory segments of doublets were determined by the following data pre-processing: first, we detected frames where tracking did not detect the loci. For up to three consecutive failures we interpolated the coordinates from before and after the failures, otherwise runs were segmented. Second, in some cases the tracking coordinates had unusually large jumps (>600 nm). When a trajectory jumped back near the prior coordinates right after a large jump we considered this jump an outlier and interpolated coordinates between before and after the jump, otherwise runs were segmented.

To compute mean square displacements, we collected all runs of doublets in the pre-processed data. Only doublet runs of a minimal length of 10 s (21 frames) were used. Within such runs, both sister loci were used to compute RMSDs. For each sister locus, in a given run squared displacements (SD) for a given time shift between two frames dt were computed as: SD(dt) = (xi+dt - xi)2 + (yi+dt - yi)2, where x and y are estimated centre positions of a single sister locus at i and i+dt time points. Squared displacements were accumulated for all valid starting frames i=1…L-dt and across both sister loci across all doublet runs across all recorded movies. Finally, RMSD(dt) were obtained by dividing the accumulated sum(SD(dt)) by the number of SD(dt) for the given dt. We show RMSDs only up to 50 s, since larger times are dominated by a few or only one long doublet run.

Anomalous diffusion coefficients D and exponents α in the relation RMSD(dt) = D*tα/2 were obtained by linear fits of log10(RMSD) versus log10(dt) plots. For the linear fits, RMSDs from t_min_fit=0.5 s to t_max_fit=10s were used.

### Sample numbers

The split kinetics of the labelled locus was characterized by imaging untreated (Fig. 2, Fig. 6, Fig. S3) or Sororin depleted and control (Fig. 8, Fig. S4) cells over the cell cycle every 36 minutes. For each locus, these experiments were repeated three times (n ≥ 5 cells per experiment), except untreated Chr1q22 and Chr3q29 and Sororin depleted Chr1q22, Chr3q29 and Chr6q25 loci were imaged 4 times and two replicas were merged into one experiment when calculating the fraction of doublets due to insufficient cell numbers. In these experiments cells were synchronized using mitotic shake-off (see above), except in one out of three replicates for untreated Chr1q22, Chr3q29, Chr4q35b, Chr6q25, Chr7q36 and Chr17p13 loci cells were growing asynchronously. In Sororin depletion experiment in 2/3 experiments for Chr1q23, in 1/3 experiments for Chr3q29 and in 1/3 experiments for Chr6q29 cells were growing asynchronously.

For G2, inter-chromatin distance measurements were pooled from 4 time points [2.4 - 0.6 h] before mitosis resulting in the following combined sample numbers from three experiments for untreated HeLa cells: Chr1p36 (n = 207), Chr1q22 (n = 288), Chr1q42 (n = 205), Chr2q14 (n = 147), Chr3q29 (n = 242), Chr4q34 (n = 201), Chr4q35a (n = 165), Chr4q35b (n = 223), Chr6q25 (n = 196), Chr7q36 (n = 243), Chr8p21 (n = 179), Chr10p13 (n = 220), Chr10q26 (n = 150), Chr11p15 (n = 232), Chr12q24 (n = 184), Chr17p13 (n = 264) and for hTERT-RPE1 cells: Chr1p36 (n = 190), Chr3q29 (n = 184), Chr4q35b (n = 204), Chr6q25 (n = 304), Chr17p13 (n = 235). From this data, a subset of inter-chromatid distances was randomly sampled for every locus to enable unbiased data visualization and comparison of different loci (n = 120 for Fig. 2 B and n = 100 for Fig. 3 E). The fraction of doublets in G2 cells was calculated in each of the 3 experiments (n ≥ 30 inter-chromatid distance measurements each) and presented as mean ± SEM (Fig. 2, D; Fig. 3 F; Fig. 4, B and D; Fig. 5, D, F and G). For Fig. 2 C, mitotic distances were measured in a single frame of untreated cells in mitosis in three experiments resulting in combined sample numbers: Chr1p36 (n = 34), Chr1q22 (n = 48), Chr1q42 (n = 38), Chr2q14 (n = 33), Chr3q29 (n = 33), Chr4q34 (n = 40), Chr4q35a (n = 31), Chr4q35b (n = 37), Chr6q25 (n = 23), Chr7q36 (n = 45), Chr8p21 (n = 31), Chr10p13 (n = 33), Chr10q26 (n = 31), Chr11p15 (n = 33), Chr12q24 (n = 42), Chr17p13 (n = 44). From this data, the fraction of doublets in mitosis was calculated from n ≥ 7 inter-chromatid distance measurements from each of the 3 experiments and presented as mean ± SEM (Fig. 2 E).

For S phase, inter-chromatin distance measurements were pooled from 6 time points [9 - 6 h] before mitosis resulting in the following combined sample numbers from all experiments: Chr1q22 (ncontrol = 345; nsiSororin = 294), Chr3q29 (ncontrol = 212; nsiSororin = 220), Chr6q25 (ncontrol = 285; nsiSororin = 575), Chr17p13 (ncontrol = 292; nsiSororin = 254). The fraction of doublets in S phase was calculated from n ≥ 35 inter-chromatid distance measurements per condition per locus per experiment and presented as mean ± SEM (Fig. 8 F).

The fraction of doublets through the entire cell cycle was calculated from ≥ 10 inter-chromatid distance measurements in a sliding window of three time points (except mitosis, t = 0 h) and presented as mean ± SEM at each time point (middle of the sliding window) (Fig. 6, D; Fig. 7 A; Fig. 8 E; Fig. S3, Fig. S4, D, F and H). Cumulative histograms of the first detected doublet were calculated from the longest and most complete trajectories pooled from 3 experiments: n = 40 trajectories (Fig. 6 D; Fig. 7 B; Fig. S3) and n = 30 trajectories (Fig. S4, E, G and I). 20 longest and most complete trajectories from three experiments are shown in Fig. 6 C and Fig. S3. The fraction of doublets after the initial split was calculated from the data as in Fig. 6 C and D and Fig. S3 from at least 5 measurements per time point per experiment in 3 independent experiments and presented as mean ± SEM in Fig. 6 E and from at least 10 measurements per time point merged from 3 experiments in Fig. 7 D.

In Fig. 6 F and G, and Fig. 8 G an H, measurements from indicated time points from 3 experiments were merged together: 6 time points for [9 - 6 h], 5 time points for [5.4 – 3 h] and 4 time points for G2 [2.4 – 0.6 h]. In Fig. 6 F and G, sample numbers for individual cell cycle stages are as follows: [9 - 6 h] n = 302 (nsinglet = 222, ndoublet = 80), [5.4 - 3 h] n = 311 (nsinglet = 187, ndoublet = 124), G2 n = 260 (nsinglet = 141, ndoublet = 119). In Fig. 8 G, sample numbers for individual cell cycle stages are as follows: for control treatment in [9 - 6 h] n = 292 (nsinglet = 191, ndoublet = 101), in [5.4 - 3 h] n = 242 (nsinglet = 127, ndoublet = 115), in G2 n = 194 (nsinglet = 75, ndoublet = 119); for Sororin depleted cells in [9 - 6 h] n = 253 (nsinglet = 102, ndoublet = 151), in [5.4 - 3 h] n = 188 (nsinglet = 61, ndoublet = 127), in G2 n = 142 (nsinglet = 31, ndoublet = 111);

In Fig. 6 I, for mean square displacement measurements cells were imaged in two independent experiments resulting in the following combined sample numbers: Chr1q22 (n = 44 trajectories from 26 cells), Chr3q29 (n = 51 trajectories from 31 cells), Chr6q25 (n = 56 trajectories from 40 cells), Chr17p13 (n = 58 trajectories from 37 cells).

For Fig. 5 C, the sample numbers for nuclear positioning measurements are as follows: Chr1p36 (n = 207), Chr1q22 (n = 288), Chr1q42 (n = 205), Chr2q14 (n = 147), Chr3q29 (n = 242), Chr4q34 (n = 201), Chr4q35a (n = 165), Chr4q35b (n = 223), Chr6q25 (n = 196), Chr7q36 (n = 243), Chr8p21 (n = 179), Chr10p13 (n = 220), Chr10q26 (n = 150), Chr11p15 (n = 232), Chr12q24 (n = 184), Chr17p13 (n = 264).

For Fig. 3, FISH was repeated in 6 independent experiments, except for Chr17p13 (n = 3), resulting in the following combined numbers of inter-chromatid measurements from all the experiments: Chr1q22 (nWT = 600, nChr1q22 = 469), Chr3q29 (nWT = 607, nChr3q29 = 505), Chr4q35b (nWT = 591, nChr4q35b = 512), Chr6q25 (nWT = 856, nChr6q25 = 799), Chr7q36 (nWT = 507, nChr7q36 = 441), Chr17p13 (nWT = 239, nChr17p13 = 326). For unbiased visualization, a subset of these data (n = 230) was randomly sampled (Fig. 3 C). The fraction of doublets was calculated in each experiment from n ≥ 34 measurements and presented as mean ± SEM in Fig. 3 D.

For Fig. S1, the numbers of labelled alleles for each locus were counted in indicated numbers of cells: Chr1p36 (n = 36), Chr1q22 (n = 31), Chr1q42 (n = 33), Chr2q14 (n = 26), Chr3q29 (n = 40), Chr4q34 (n = 39), Chr4q35a (n = 46), Chr4q35b (n = 29), Chr6q25 (n = 41), Chr7q36 (n = 10), Chr8p21 (n = 43), Chr10p13 (n = 42), Chr10q26 (n = 36), Chr11p15 (n = 44), Chr12q24 (n = 40), Chr17p13 (n = 25).

For Fig. S2 E, the numbers of cells were counted at time point 0 h and in 24 h in three independent experiments resulting in the following combined numbers of cells: wild type HeLa Kyoto (n0h = 742, n24h = 1655), dCas9-mEGFP parental cell line (n0h = 657, n24h = 1575), Chr1q22 (n0h = 750, n24h = 1741), Chr1q4 (n0h = 466, n24h = 1079), Chr4q35b (n0h = 601, n24h = 1419), Chr6q25 (n0h = 680, n24h = 1502), Chr7q36 (n0h = 567, n24h = 1219), Chr10p13 (n0h = 660, n24h = 1534), Chr10q26 (n0h = 508, n24h = 1236), Chr17p13 (n0h = 558, n24h = 1280). For Fig. S2 F, n = 3 experiments, 30 cells each. For Fig. S2 O, ndCas9-mEGFP = 26 cells and ndCas9-SunTag = 25 cells resulting in combined number of labelled loci quantified: ndCas9- mEGFP = 76 and ndCas9-SunTag = 75. For Fig. S2 L, n = 10800 images were simulated.

For Fig. S4 B, the number of Chr17p13 cells arrested in mitosis was calculated from 3 independent experiments resulting in combined numbers of n = 98 cells in control and n = 110 cells in siSororin treatment. For Fig. S4 C, the cumulative histogram of the cell cycle duration is calculated for Chr17p13 locus from three experiments with the following combined cell numbers: ncontrol = 137, nsiSororin = 167.

### Statistical analysis

All statistical tests were performed with R 3.1.2 using stats and lawstat packages. The normality of data distribution was tested using Kolmogorov-Smirnov test and the variance was tested using Levene test.

For two groups of normally distributed data, two-sided unpaired Welch’s t-test was performed: in Fig. 5 E (p = 0.02); in Fig. 5 F (p = 0.02); in Fig. 5 G (p = 0.10); in Fig. 5 H (p = 0.01); in Fig. 5 I (p = 0.10); in Fig. 5 J (p = 0.17); in Fig. 5 K (p = 0.07); in Fig. 5 L (p = 0.02); in Fig. 5 M (p = 3.0*10-3); in Fig. 7 H: Chr1q22, (p = 0.02), Chr3q29 (p = 0.02), Chr6q25 (p = 0.03), Chr17p13 (p = 0.01); in Fig. 7 J Chr1q22 (p = 0.11), Chr3q29 (p = 0.06), Chr6q25 (p = 0.76), Chr17p13 (p = 0.09); in Fig. S2 O (p = 4.2*10-10); in Fig. S4 B (p = 0.01). Three and more groups of normally distributed data were tested with one way ANOVA not assuming equal variance: in Fig. 2 D (p = 6.8*10-8); in Fig. 2 E (p = 0.75); in Fig. 3 D (pwt = 3.9*10-8); in Fig. S2 E (p = 0.12); in Fig. S2 F (p = 0.97);

Two groups of not-normally distributed data were compared using two-sided unpaired Wilcoxon rank-sum test: in Fig. 3 C: Chr1q22 (p = 0.80), Chr3q29 (p = 0.23), Chr4q35b (p = 0.85), Chr6q25 (p = 0.33), Chr7q36 (p = 0.12), Chr17p13 (p = 0.33); in Fig. 7 G: Chr1q22 (p < 2.2*10-16), Chr3q29 (p = 1.3*10-13), Chr6q25 (p = 2.4*10-4), Chr17p13 (p = 3.9*10-11); in Fig. 7 I: Chr1q22 (p = 3.4*10-11), Chr3q29 (p = 6.5*10-8), Chr6q25 (p = 0.03), Chr17p13 (p = 1.1*10-7). More than three groups of not normally distributed data were compared using Kruskal-Wallis rank sum test: in Fig. 5 C (p < 2.2 * 10-16).

### Data plotting

All data was plotted using R 3.12 using graphics and beeswarm packages except for Fig. 6, F-I and Fig. 8, G and H, data was plotted using Matlab R2017b. All microscopy images for the figures were processed with Fiji, using Gaussian blur denoising (0.8 pixel diameter) and cropping or projections as indicated. ChIP-sequencing tracks for SMC3 and Sororin were visualized with integrated genome viewer (IGV 2.3.66) (Robinson et al., 2011).

---

**Figure S1.**
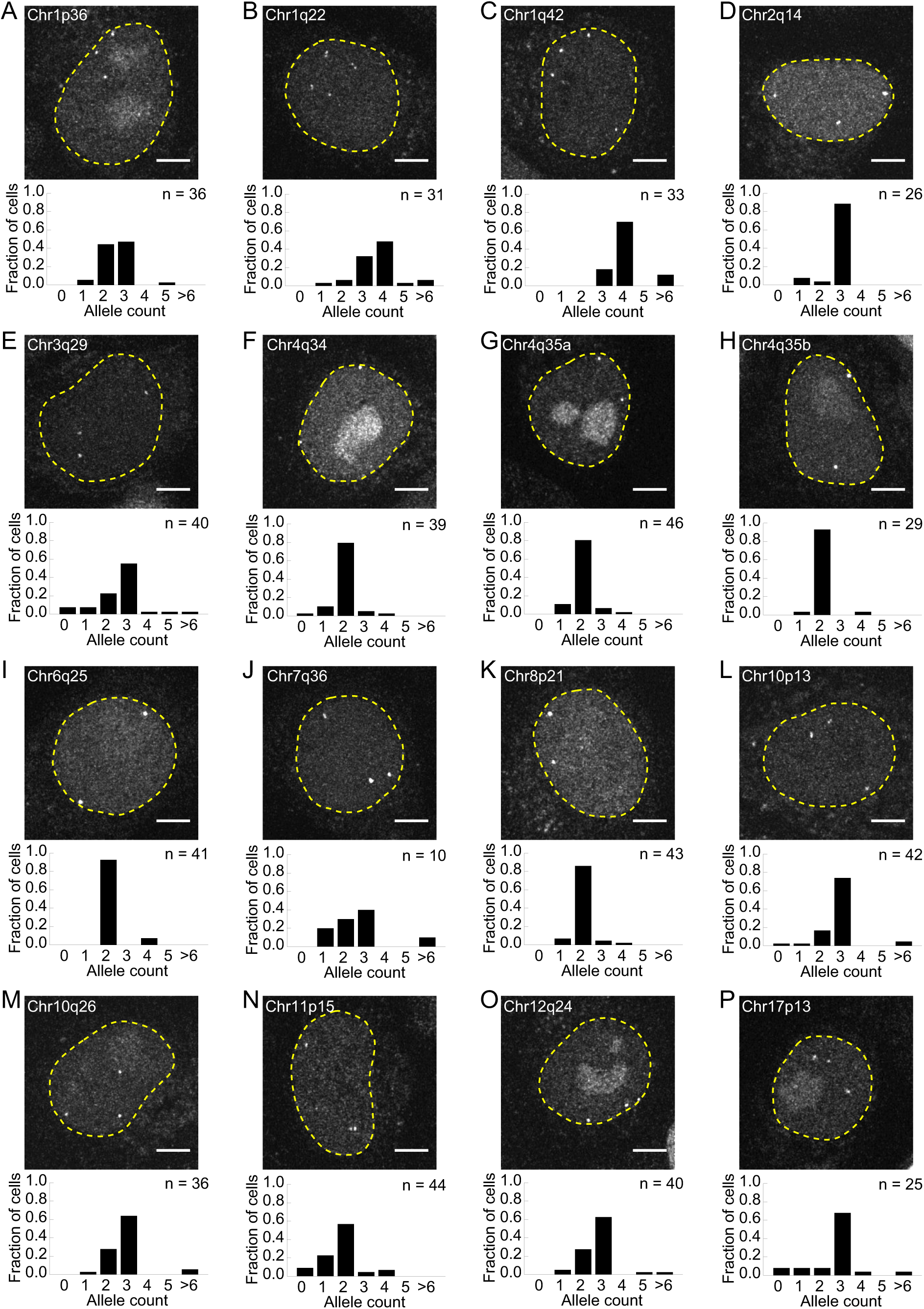
Allele counts of dCas9-mEGFP labelled foci per cell. (A-P) Confocal images of representative live cells for each of the 16 cell lines with dCas9-mEGFP/sgRNA-labelled loci. Images show projections of 21 z-sections; dashed lines indicate nuclear rim. Bar-graphs show the distribution of fluorescent dot numbers in each cell line. Fluorescent dots were counted in S or G2 cells and pairs of foci that were closer than 2 μm were scored as a doublet representing a single replicated locus; n = number of cells. Scale bars, 5 μm.

**Figure S2.**
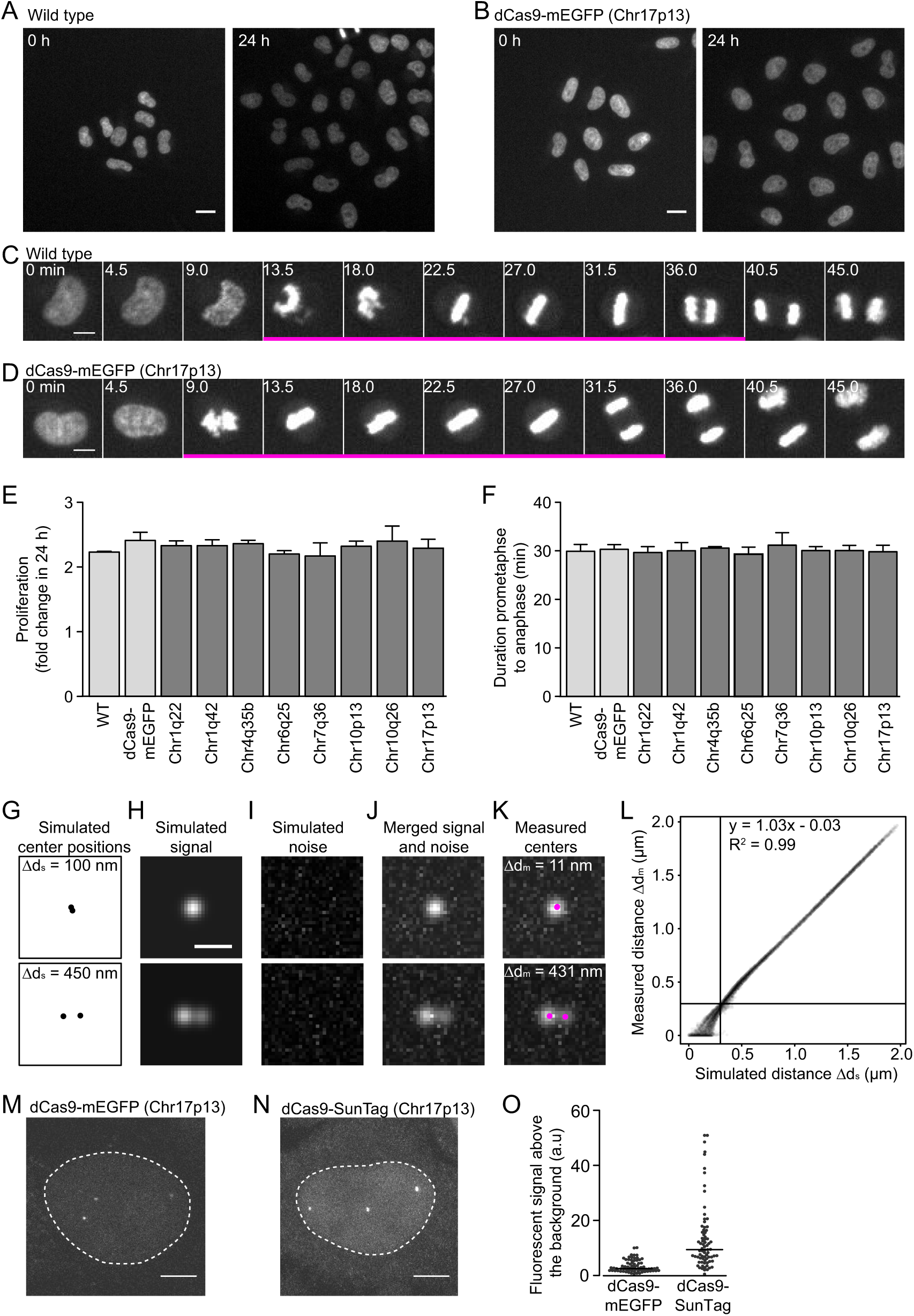
Characterisation of dCas9-mEGFP/sgRNA cell lines and validation of image analysis procedure. (A-B) Time-lapse images of wild type cells (A) or cell line expressing dCas9-mEGFP and sgRNA targeting Chr17p13 (B) were stained with SiR-Hoechst. (C-D) Example of a wild type cell (C) and Chr17p13-labelled cell (D) progressing through mitosis. Single z-sections, numbers indicate time (min), magenta bar indicates prometaphase to anaphase frames. (E) Quantification of cell proliferation from images as in A and B for cell lines as indicated. Differences are not significant (p = 0.12 by one way ANOVA test). Error bars indicate mean ± SD; n = 3 experiments. (F) Quantification of mitotic duration (prometaphase until anaphase) in images as in C and D. Error bars indicate mean ± SD; n = 3 experiments, 10 cells each. Differences are not significant (p = 0.97 by one way ANOVA test). (G-L). The accuracy of inter-chromatid distance measurements by the 2D Gaussian mixture model fitting procedure (Fig. 1, D-H) was estimated from simulated images. (G) Two simulated point light sources at random positions with simulated distance Δds = 100 nm (upper panel) and Δds = 450 nm (lower panel). (H) Two 2D Gaussian functions approximate signal in the microscopy images (H) around the simulated point light sources in (G). Variance and amplitude of the Gaussian functions are randomly sampled from the respective distributions observed in real microscopy images of Chr3q29 locus. (I) Image noise is derived by randomly sampling noise values for each pixel from the distribution of noise intensities observed in real microscopy images of Chr3q29 locus. (J) Simulated images resulting from merged signal in (G) and noise in (H). (K) Centre position (magenta) estimated in simulated images as in (J) by fitting mixture of two 2D Gaussian functions as in (Fig. 1, D-H). Δdm indicate measured distances between measured centre positions. (L) Relationship between simulated distances Δds in simulated images and distances Δdm measured in these images using mixture of two 2D Gaussian function fitting procedure as in (Fig. 1, D-H). Vertical and horizontal lines indicate 300 nm threshold. n = 10800 simulated images. (M-O) Comparison of signal intensity of dCas9-mEGFP and dCas9-SunTag labelled loci. (M-N) Example cell nuclei (dashed line) with Chr17p13 locus labelled using dCas9-mEGFP (M) and dCas9-SunTag (N).(O) Signal intensity above the nucleoplasmic background at Chr17p13 locus labelled with dCas9-mEGFP and dCas9-SunTag in the corresponding stable cell lines (p = 4.2*10-10 by unpaired two-sided t test). Scale bars, 20 μm (A), 5 μm (C-D and M-N), 1 μm (H).

**Figure S3.**
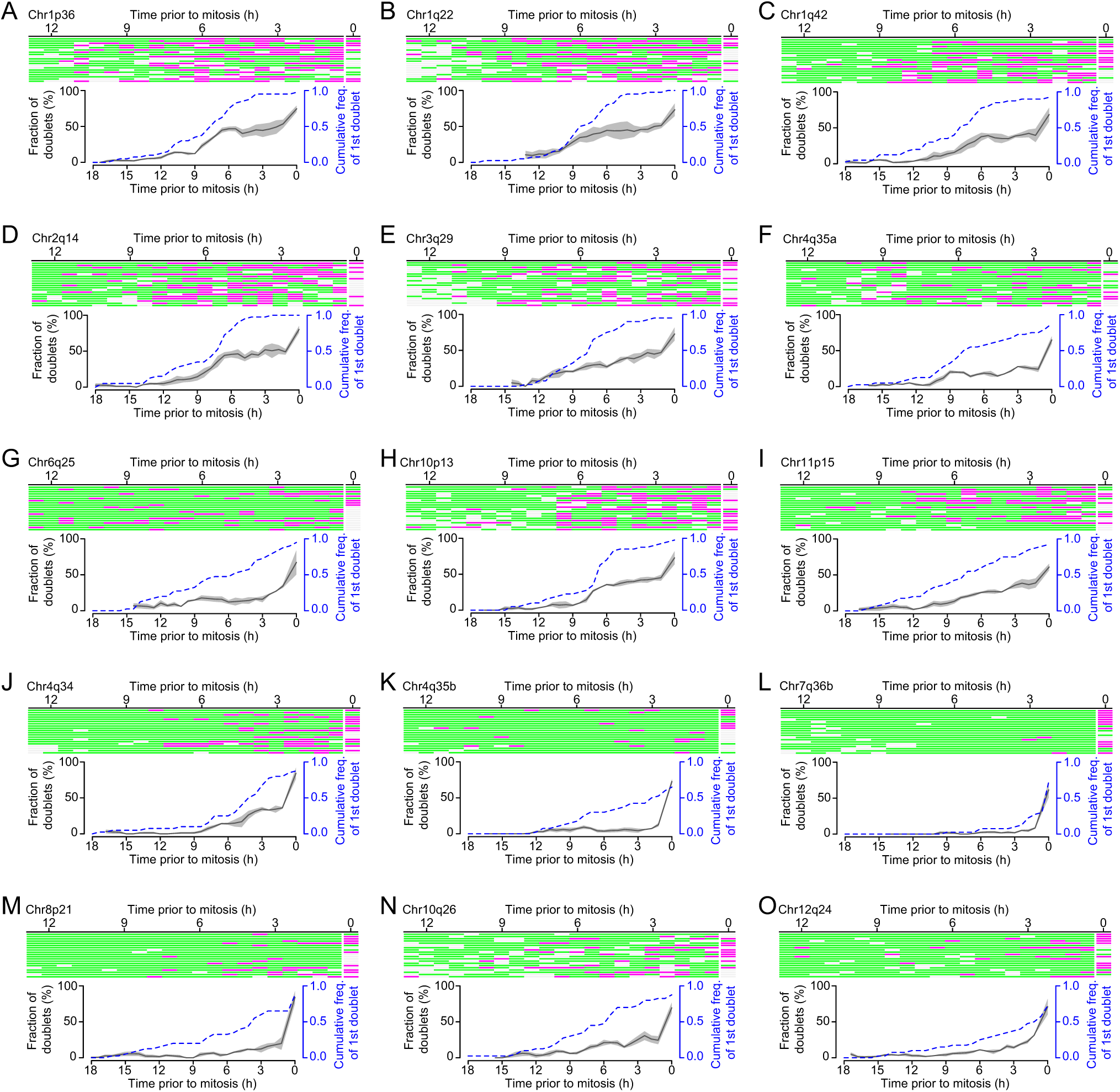
Kinetics of sister locus resolution from S-phase until mitosis. Full dataset for individual locus trajectories as shown for Chr17p13 in Fig. 6 B-E, for the other 15 loci. Upper panels show individual allele trajectories during cell cycle progression. Doublets (magenta), singlets (green) and missing data points (light grey) were automatically annotated based on Gaussian mixture model fitting as in Fig. 1 D-H (n = 20 trajectories for each genomic site). Lower panels indicate fraction of doublets (solid line and shaded area indicate mean ± SEM, respectively; n = 3 experiments). Dashed lines indicate cumulative frequency of the first detected doublet, n = 40 trajectories. (A-I) Loci replicating during early S phase, as shown in Fig. 6 F and G. (J-O) Loci replicating during late S phase, as shown in (Fig. 6, H and I).

**Figure S4.**
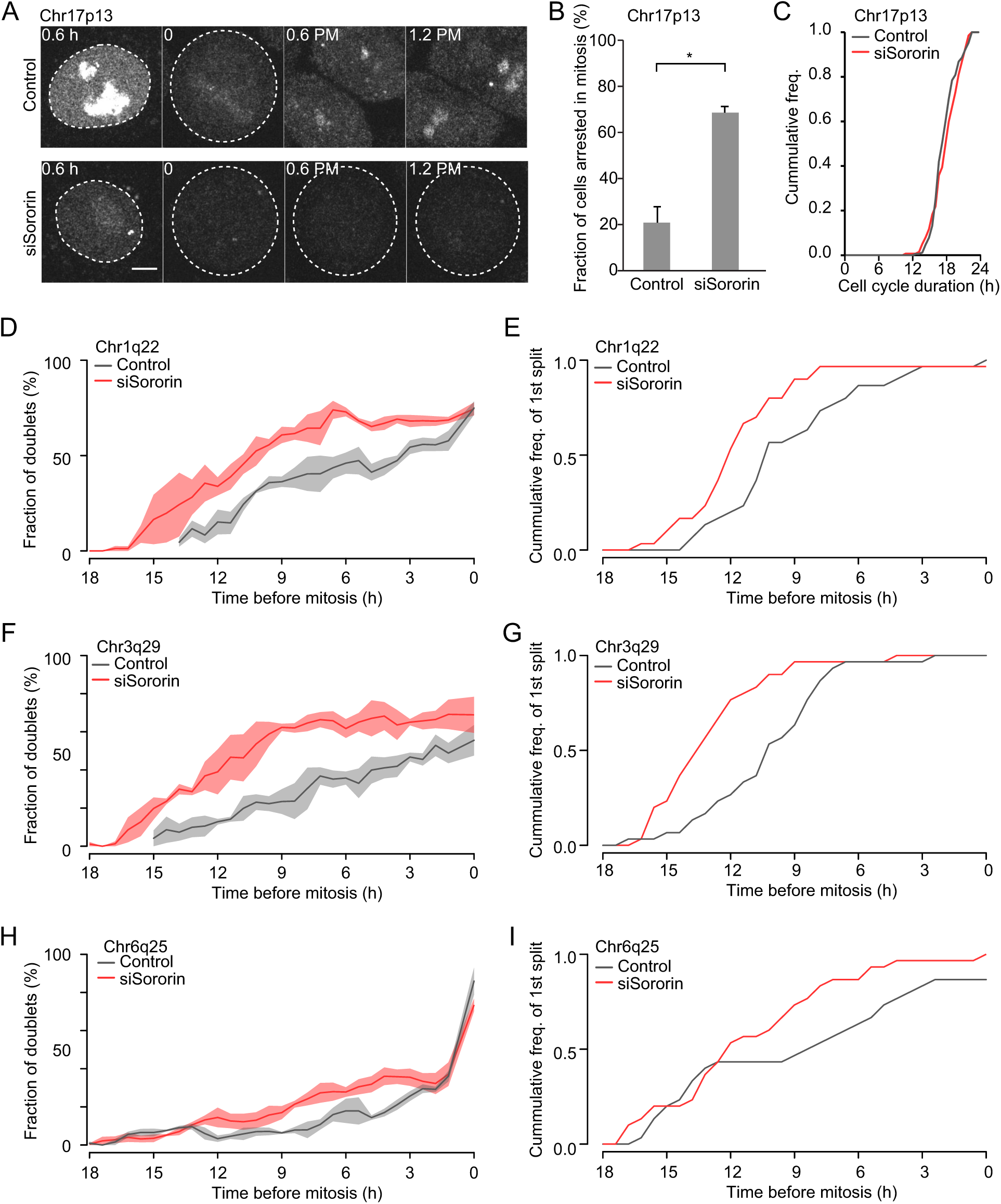
Local sister chromatid cohesion persists for several hours after DNA replication. (A) Example images of a control and Sororin depleted cell entering mitosis. As judged by rounded appearance, the control cell stays in mitosis only for 1 frame, thus mitosis takes less than 1h 12 min. The Sororin depleted cell, on the other hand, arrests in mitosis for more than 3 frames, thus mitosis takes longer than 1 h 12 min after Sororin depletion. Maximum intensity projections of z-sections 3.5 and 10.5 μm above the cover glass, corresponding to the middle section of interphase and mitotic cell respectively. Scale bar 5 μm. (B) Cells arrest in mitosis after Sororin depletion, quantified from images as in (A). Errors represent mean ± SEM, n = 3 experiments, p < 0.001 derived from two-sided unpaired t test. (C) Cumulative histogram of cell cycle duration from three experiments of cells with Chr17p13 locus labelled in control (n = 98) and Sororin depletion (n = 110), as in Fig. 8, A-F. Cells were synchronized by mitotic shake off and imaged while progressing through cell cycle as in Fig. 1. Cell cycle duration was measured as time from the beginning of imaging to the first frame in mitosis (rounding of the cell as in Fig. S4 A). (D, F, H) The fraction of doublets over the cell cycle in control (grey) and Sororin depleted cells (red) at (D) Chr1q22, (F) Chr3q29 and (H) Chr6q25 loci. Shaded area indicates mean ± SEM; n = 3 experiments. (E, G, I) Cumulative frequency of the first observed doublets in control (grey) and Sororin depleted (red) cells at (E) Chr1q22, (G) Chr3q29 and (I) Chr6q25 loci; n = 30 trajectories.

**Supplementary Table 1.**
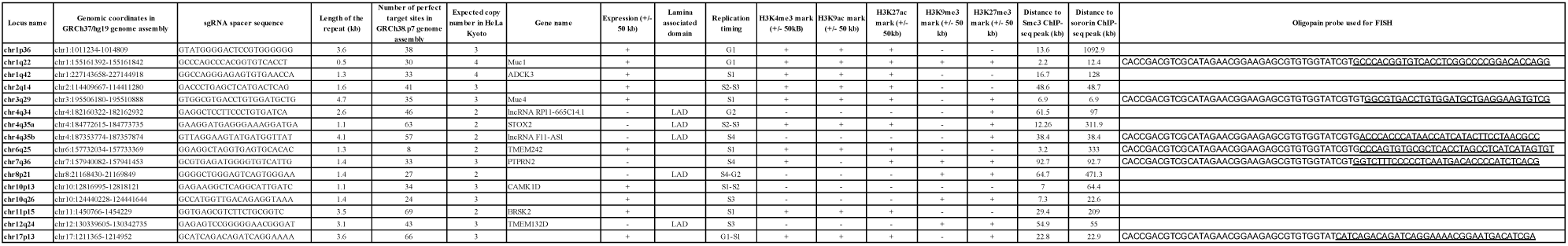
Positions and genomic neighbourhoods of dCas9-mEGFP/sgRNA-labelled endogenous loci. Genomic coordinates from GRCh37/hg19 assembly; copy number for HeLa Kyoto from (Landry et al., 2013); perfect targets in GRCh38.p7 genome assembly; expression data from LAD localization from (Guelen et al., 2008); replication timing, gene expression, and histone modification data for HeLa from (Hansen et al., 2010; ENCODE Project Consortium, 2012); SMC3 and Sororin enrichment sites from (Ladurner et al., 2016).

